# Serotonergic Neurons Translate Taste Detection into Internal Nutrient Regulation

**DOI:** 10.1101/2021.05.13.444014

**Authors:** Zepeng Yao, Kristin Scott

## Abstract

The nervous and endocrine systems coordinately monitor and regulate nutrient availability to maintain energy homeostasis. Sensory detection of food regulates internal nutrient availability in a manner that anticipates food intake, but sensory pathways that promote anticipatory physiological changes remain unclear. Here, we identify serotonergic (5-HT) neurons as critical mediators that transform gustatory detection by sensory neurons into the activation of insulin-producing cells and enteric neurons in *Drosophila*. One class of 5-HT neurons responds to gustatory detection of sugars, excites insulin-producing cells and limits consumption, suggesting that they anticipate increased nutrient levels and prevent overconsumption. A second class of 5-HT neurons responds to gustatory detection of bitter compounds and activates enteric neurons to promote gastric motility, likely to stimulate digestion and increase circulating nutrients when food quality is poor. These studies demonstrate that 5-HT neurons relay acute gustatory detection to divergent pathways for longer-term stabilization of circulating nutrients.

## Introduction

Animals have evolved elaborate mechanisms that engage the nervous, endocrine, and digestive systems to monitor and regulate nutrient availability. Sensory systems, including the gustatory system, identify potential food sources prior to nutrient ingestion. Endocrine and digestive systems support long-term stabilization of circulating nutrients that sustain tissue and organ function. These multiple systems interact to coordinately regulate nutrient availability and maintain energy homeostasis, often across different time scales.

Seminal studies have revealed that sensory detection of food sets in motion anticipatory physiological state changes (Chen and Knight, 2016; Power and Schulkin, 2008; Zafra et al., 2006). For instance, the detection of food by the visual, olfactory or gustatory systems influences salivation, digestive enzyme secretion and hormone release. These sensory-evoked responses are called cephalic phase responses, as they are elicited by the brain prior to nutritive state changes (Power and Schulkin, 2008; Zafra et al., 2006). One illustrating example is cephalic phase insulin release in mammals. It has been known for decades that sensory detection of food triggers an initial insulin pulse that occurs before nutrients are absorbed, serving to increase glucose storage prior to blood glucose increases (Power and Schulkin, 2008; Teff, 2000; Zafra et al., 2006). Another example is anticipatory regulation of feeding-related neurons in the arcuate nucleus of the mouse hypothalamus: AgRP (Agouti-Related Peptide) neurons are activated by hunger signals and this hunger-related activity rapidly decreases in response to the smell or sight of food alone, in the absence of ingestion (Betley et al., 2015; Chen and Knight, 2016; Chen et al., 2015; Mandelblat-Cerf et al., 2015). Similarly, the activity of thirst neurons in the subfornical organ is rapidly downregulated upon initiating water ingestion, prior to blood osmolality changes (Augustine et al., 2018; Zimmerman et al., 2016). A better understanding of how acute sensory detection elicits longer-term physiological changes will provide insight into the anticipatory regulation of nutrient homeostasis.

Taste detection is a key signal of nutrient availability that precedes physiological changes that accompany food intake. For example, gustatory detection of nutrients such as sugars precedes nutrient intake, digestion, increased circulating nutrients, and satiation. In contrast, bitter taste detection signals poor food quality, a lack of external nutrient availability, and potential nutrient deprivation in the near future. While taste detection is poised to communicate anticipated changes in nutrient status to the endocrine and digestive systems, the taste pathways that coordinate these systems remain unclear.

*Drosophila*, like mammals, monitors external nutrients using the gustatory system and regulates internal nutrient availability using the endocrine and digestive systems. Key features of these systems are shared in flies and mammals, including gustatory neurons tuned to sweet and bitter taste compounds (Yarmolinsky et al., 2009), and internal nutrient regulation via insulin and glucagon equivalents (Ahmad et al., 2020; Leopold and Perrimon, 2007). The numerical simplicity of the fly nervous system, the ability to monitor and manipulate single neurons, and the potentially short relays between the brain, the endocrine system and the digestive tract make *Drosophila* an ideal system for examining common mechanisms underlying nutrient homeostasis.

Here, we investigate communication between the *Drosophila* gustatory system and endocrine and digestive systems and identify serotoninergic (or 5-hydroxytryptamine, 5-HT) neurons as critical circuit nodes that mediate anticipatory regulation of nutritional state. Using intersectional genetic approaches, we identified and characterized two distinct classes of 5-HT neurons that translate short-term changes in taste detection into longer-term changes in physiological state. One class of 5-HT neurons is activated by sugar taste detection prior to ingestion. These neurons excite insulin-producing neurons and limit sugar consumption, suggesting that they promote insulin release in anticipation of sugar intake and prevent overconsumption. A second class of 5-HT neurons is activated by gustatory detection of bitter compounds and activates enteric neurons that promote gastric motility. As the detection of bitter compounds is often indicative of harmful or toxic compounds, these neurons likely function to stimulate food digestion to replenish circulating nutrients when food quality is poor. Together, our work reveals that gustatory stimuli activate 5-HT neurons to anticipate nutrient availability and coordinate endocrine and digestive function for longer-term stabilization of available nutrients.

## Results

### Sugar and Bitter Tastes Activate Distinct 5-HT Neurons

Serotonin powerfully modulates appetite and food intake across animals (Tecott, 2007). The *Drosophila* central brain contains ∼90 5-HT neurons (Pooryasin and Fiala, 2015), offering a tractable model to study 5-HT neurons that modulate feeding. To examine whether 5-HT neurons in *Drosophila* contribute to taste processing and feeding regulation, we first tested whether they respond to taste sensory input. The primary gustatory center of the fly brain is the subesophageal zone (SEZ) which is innervated by the axons of gustatory neurons from fly taste organs, including the proboscis, mouthparts, and legs (Stocker, 1994). Therefore, we hypothesized that 5-HT neurons in this brain region might respond to taste sensory detection.

We monitored taste-induced activity in 5-HT neurons throughout the SEZ by *in vivo* GCaMP6s calcium imaging (Chen et al., 2013; Harris et al., 2015). To selectively label 5-HT neurons, three independent *Tryptophan hydroxylase* (*Trh*) *Gal4* lines (Alekseyenko et al., 2010; Shearin et al., 2013) were used for imaging studies. The esophagus was severed to evaluate whether taste sensory stimulation activated 5-HT neurons independent of ingestion. Sucrose stimulation of proboscis gustatory neurons activated SEZ 5-HT neurons that have dense and diffuse arbors (Figures 1A-1C, upper panels). Interestingly, bitter stimulation activated different SEZ 5-HT neurons that have discrete branched arborizations (Figures 1A and 1B, lower panels). Whereas two *Trh-Gal4* lines, *Trh-Gal4(S)* and *Trh-Gal4(K3),* contained both sugar and bitter responding cells (Figures 1A and 1B), *Trh-Gal4(K2)* neurons responded only to sugar sensory stimulation (Figure 1C). Co-labelling studies revealed that *Trh-Gal4(S)* and *Trh-Gal4(K3)* labelled all SEZ 5-HT neurons (Figures 1D and 1E). *Trh-Gal4(K2)* labeled all SEZ 5-HT neurons except for two 5-HT neural pairs in the lateral SEZ, suggesting that the unlabeled neurons are the bitter-responding cells (Figure 1F, yellow arrows). These results demonstrate that there are two distinct classes of 5-HT neurons in the SEZ that respond to sugar and bitter taste detection, respectively.

**Figure 1.**
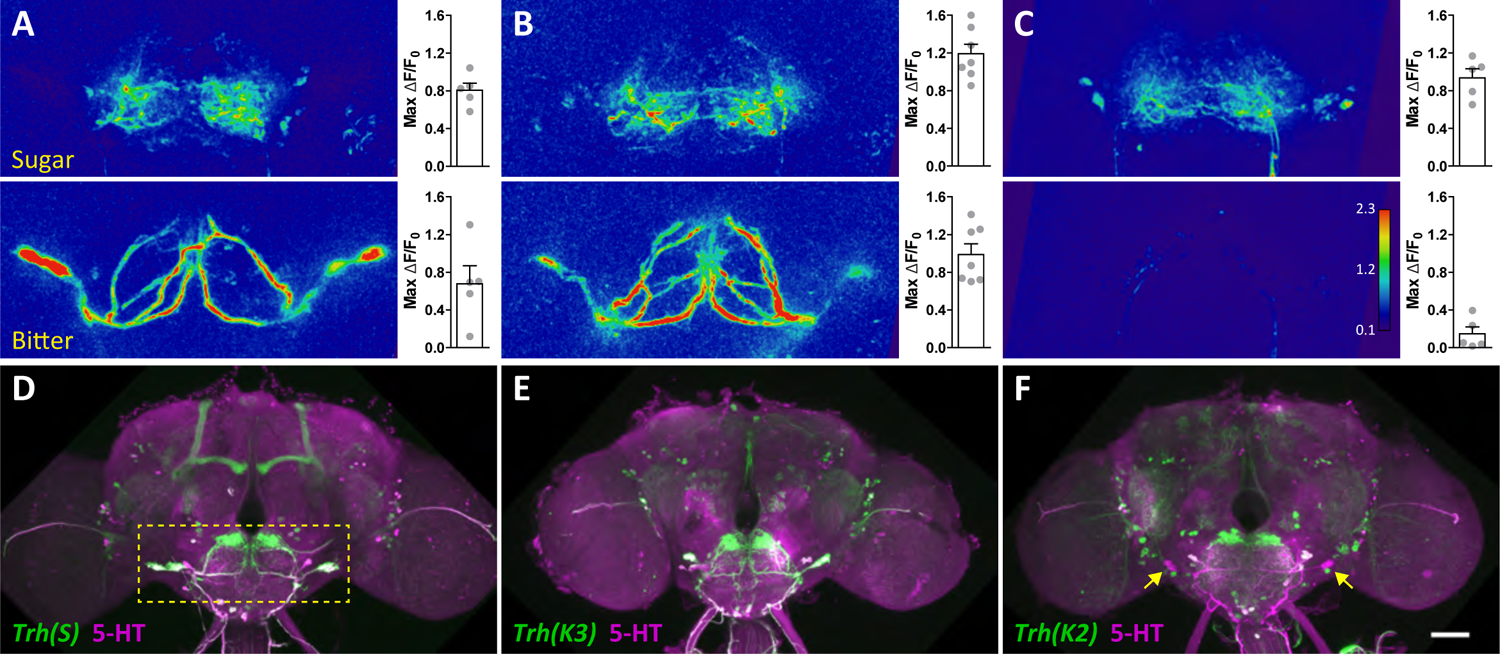
Sugar and Bitter Activate Different SEZ 5-HT neurons. (A-C) Left: Representative ΔF/F_0_ images of GCaMP6s imaging showing SEZ 5-HT neurons responding to sugar (upper panels) and bitter (lower panels) proboscis taste detection. GCaMP6s was expressed using three independent *Trh-Gal4* (*Trh: tryptophan hydroxylase*) lines characterized in (D-F). Right: Maximum ΔF/F_0_ changes for individual flies (dots) and bar plot overlay, mean ± SEM. n = 5-7 flies/genotype. (D-F) The expression patterns of *Trh-Gal4* lines used for GCaMP6s imaging in (A-C) were characterized using *UAS-CD8-GFP* reporter expression (green). Antibody staining against 5-HT is shown in magenta. Yellow dashed rectangle in (D) shows the approximate area for calcium imaging in (A-C). Yellow arrows in (F) point out two pairs of SEZ 5-HT neurons that are not labelled by *Trh-Gal4(K2)*. Scale bar, 50 μm.

### Sugar-SELs Respond to Proboscis Sugar Detection

The taste-responsive 5-HT cells belong to the anatomically defined lateral subesophageal ganglion (SEL) 5-HT neural cluster (Pooryasin and Fiala, 2015); therefore, we named the sugar and bitter responding neurons as sugar-SELs and bitter-SELs, respectively. We used an intersectional genetic approach to gain specific genetic access to the sugar-SELs to study their function. The intersection between *Trh-Gal4(K2*), expressed in sugar-SELs but not bitter-SELs (Figures 1C and 1F), and the Hox gene based *Dfd-LexA,* expressed in dorsal SEZ cells (Simpson, 2016), was used to restrict *Gal4* expression specifically to the sugar-SELs (Figure 2A; Video S1; STAR Methods).

**Figure 2.**
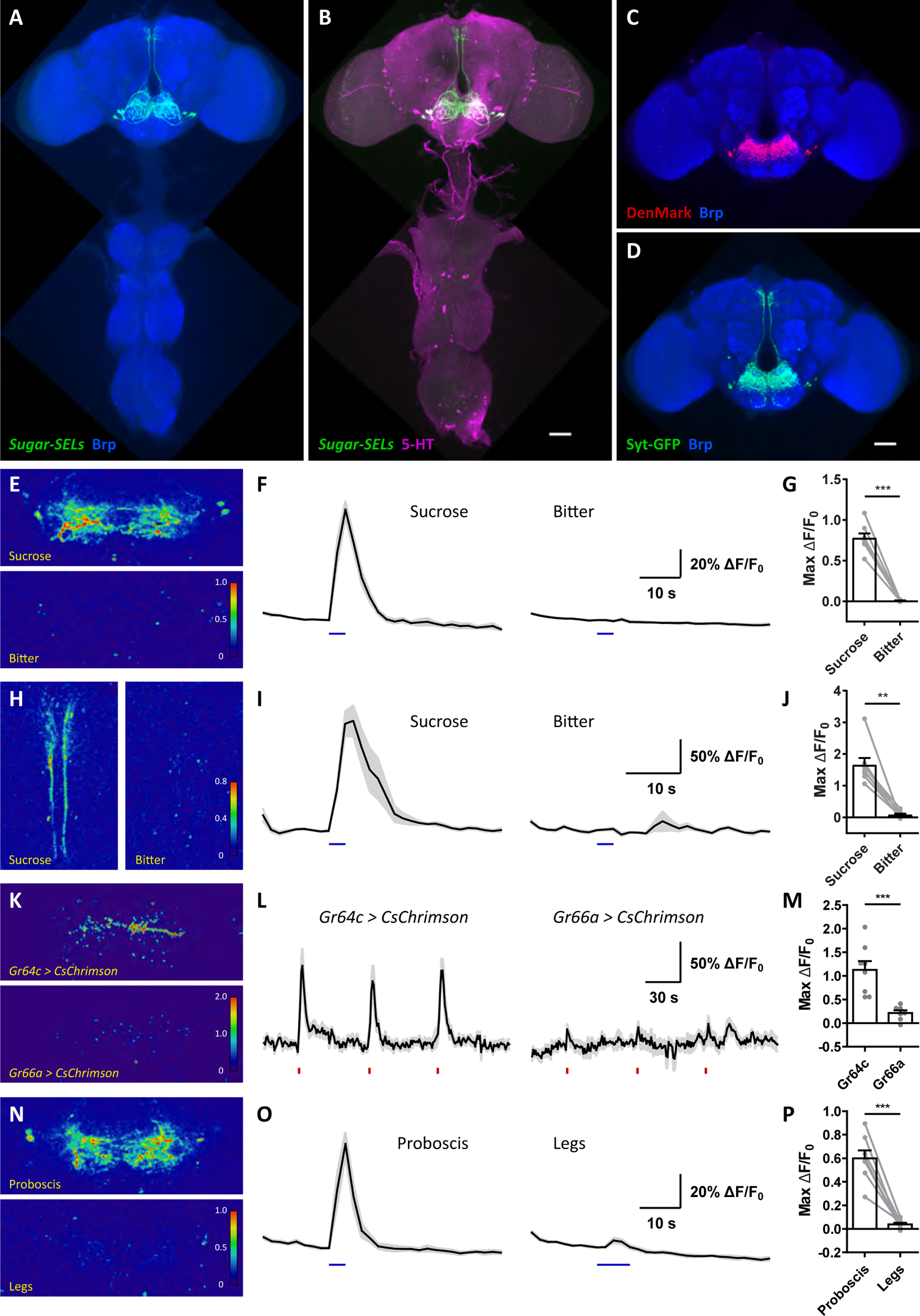
Sugar-SELs Specifically Respond to Proboscis Sugar Detection. (A and B) Sugar-SELs are labelled by the genetic intersection between *Trh-Gal4(K2)* and *Dfd-LexA* (green). Anti-Brp staining (A) shows brain and ventral nerve cord (VNC) neuropil (blue). Anti-5-HT staining (B) shows 5-HT neurons (magenta). Scale bar, 50 μm. (C and D) Dendritic compartments (C, DenMark, red) and axonal compartments (D, Syt-GFP, green) of sugar-SELs. Anti-Brp staining shows brain neuropil (blue). Scale bar, 50 μm. (E-G) Sugar-SEL arbors in SEZ respond to proboscis sucrose but not bitter detection. (H-J) Sugar-SEL projections along the median bundle respond to proboscis sucrose but not bitter detection. (K-M) Sugar-SEL PNs respond to CsChrimson-mediated optogenetic excitation of sugar (*Gr64c^+^*) but not bitter (*Gr66a^+^*) GRNs. (N-P) Sugar-SELs respond to sucrose detection on the proboscis but not on the legs. Images in (E), (H), (K), and (N) are representative ΔF/F_0_ images of GCaMP6s responses. Plots in (F), (I), (L), and (O) show mean ΔF/F_0_ traces (black lines) ± SEM (gray shading). Blue bars in (F) and (I) indicate sucrose (left) or bitter (right) stimulation. Red bars in (L) indicate red light stimulations. Blue bars in (O) indicate sucrose stimulation on the proboscis (left) or legs (right). (G), (J), (M), and (P) are scatter plots of maximum ΔF/F_0_ changes, with bar plot overlay, mean ± SEM. (G), (J), and (P): n = 8 flies; paired t-test, ** p < 0.01, *** p < 0.001. (M): n = 8 *Gr64c* flies, n = 7 *Gr66a* flies; Mann Whitney test, *** p < 0.001. See Video S1 for sugar-SEL anatomy. See Figure S1 for additional analysis of sugar-SEL subtypes.

Sugar-SELs consist of three neural pairs in the SEL cluster (Figures 2A and 2B). Sugar-SELs contain both dendritic (marked by DenMark (Nicolai et al., 2010)) and axonal (marked by Syt-GFP (Zhang et al., 2002)) arborizations in the SEZ (Figures 2C and 2D), consistent with a role in taste processing. In addition, sugar-SELs send axonal projections along the median bundle to the pars intercerebralis (PI) (Figure 2D), a neuroendocrine center. Single-cell labeling approaches (Hampel et al., 2011) revealed three different sugar-SEL morphological subtypes: an ipsilateral projection neuron (PN) (Figure S1A), a contralateral PN (Figure S1B), and a SEZ local neuron (LN) (Figure S1C). A *split-Gal4* collection designed to label individual cell types in the SEZ contained two *split-Gal4* lines that target the sugar-SEL PNs and the sugar-SEL LNs, respectively (Figures S1D-S1E’) (Sterne et al., in preparation). Sugar-SEL PNs have dendritic arbors in the SEZ and axonal arbors in the SEZ and PI (Figures S1F and S1F’). Sugar-SEL LNs have dendrites and axons in the SEZ (Figure S1G).

We confirmed that sugar-SELs respond to stimulation of the proboscis with sugar but not with bitter compounds (Figures 2E-2J). Calcium responses to sugar stimulation were detected in SEZ arbors (Figures 2E-2G) and median bundle projections (Figures 2H-2J), using flies with GCaMP6s expressed in all sugar-SELs. By using specific *split-Gal4* lines that distinguish the sugar-SEL PNs and LNs, we found that both cell populations responded to sugar stimulation (Figures S1H-S1K). To confirm that the sugar-SELs are activated by gustatory detection, we optogenetically activated sugar (*Gr64c^+^*) or bitter (*Gr66a^+^*) gustatory receptor neurons (GRNs) (Freeman and Dahanukar, 2015) (Figures S1L and S1M) using the light-gated cation channel CsChrimson (Klapoetke et al., 2014) and monitored sugar-SEL PN responses. As expected, sugar-SEL PNs consistently responded to optogenetic excitation of sugar GRNs but not bitter GRNs (Figures 2K-M). As flies have GRNs not only on the proboscis but also on other peripheral appendages such as the legs, we tested whether the sugar-SEL taste response is organ specific. Interestingly, sugar-SELs specifically responded to sugar stimulation of the proboscis but not the legs (Figures 2N-2P), suggesting that sugar-SEL function may be related to feeding rather than to food searching.

### Sugar-SELs Excite Insulin-Producing Cells

Sugar-SEL PNs send axons to the PI (Figures S1A, S1B, S1D, and S1F’), raising the intriguing possibility that they regulate neuroendocrine function. The PI contains many neurosecretory cells, including insulin-producing cells. These cells produce and secrete *Drosophila* insulin-like peptides (DILPs) and regulate sugar metabolism, similar to mammalian pancreatic β-cells (Ahmad et al., 2020; Leopold and Perrimon, 2007). We examined whether sugar-SEL PNs arborize in proximity to Dilp2 insulin-producing cells, and found that the neurites of the two cell types overlap extensively in the SEZ, the median bundle, and the PI (Figures 3A-3C and S2A), suggestive of potential synaptic connections.

**Figure 3.**
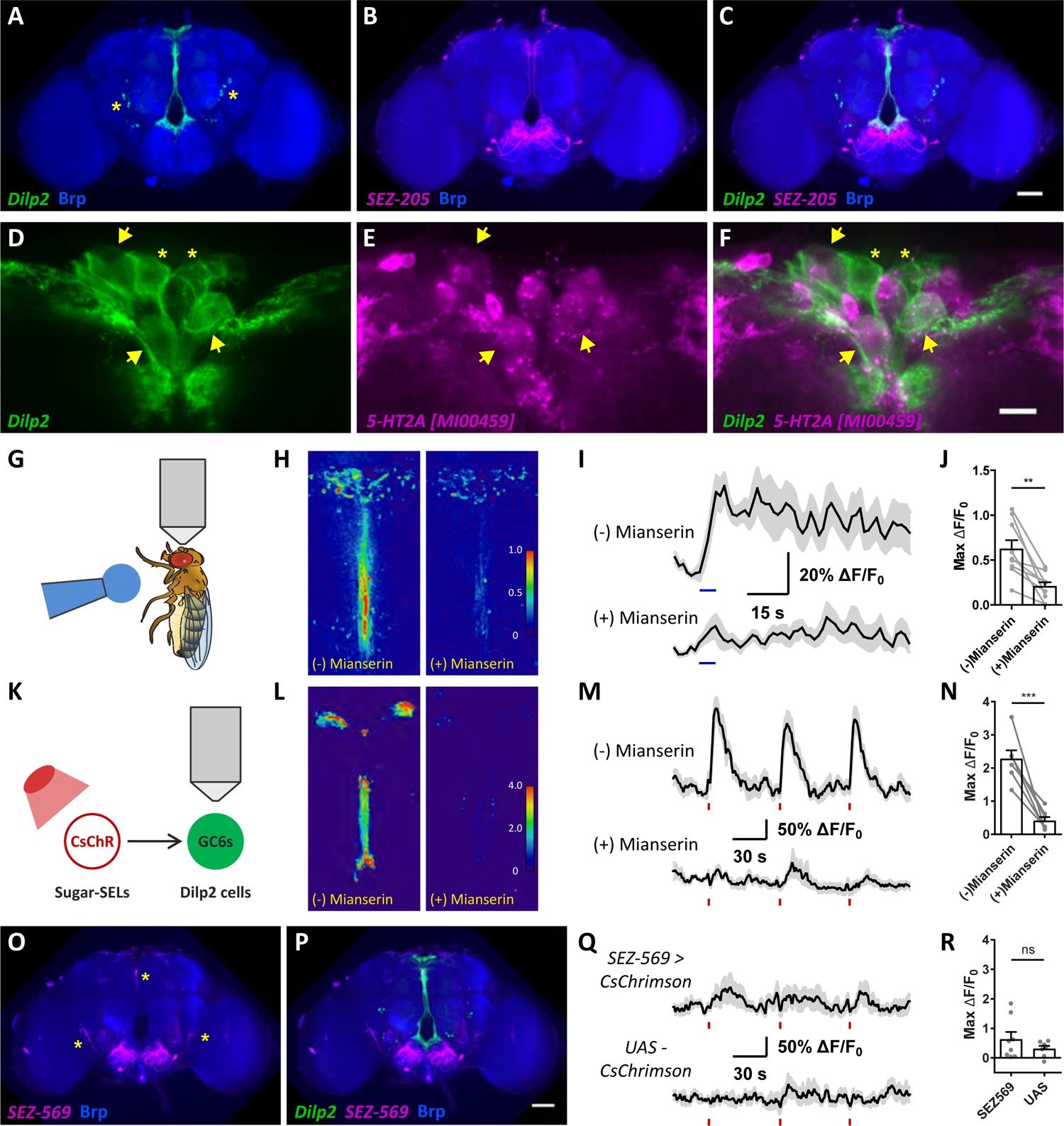
Sugar-SEL PNs Excite Insulin-Producing Cells, Likely Through 5-HT2A. (A-C) Anatomical proximity of Dilp2 insulin-producing cells (*Dilp2-LexA>LexAop-CD8-GFP*, green) and sugar-SEL PNs (*SEZ-205>UAS-CD8-RFP*, magenta). Anti-Brp staining shows brain neuropil (blue). Yellow asterisks (A) mark non-specific cells labelled by *Dilp2-LexA*. Scale bar, 50 μm. (D-F) A subset of Dilp2 cells expresses 5-HT2A. Dilp2 cells are labeled by *Dilp2-LexA>LexAop-CD8-GFP* and stained for GFP (green). 5-HT2A expressing cells are labeled by *5-HT2A[MI00459]-Gal4>UAS-CD8-RFP* and stained for RFP (magenta). Yellow arrows mark Dilp2 cells that express 5-HT2A; Yellow asterisks mark Dilp2 cells that do not express 5-HT2A. Scale bar, 10 μm. (G-J) Dilp2 cells respond to proboscis sucrose detection. (G) Experiment schematic. (H) Representative GCaMP6s ΔF/F_0_ images of Dilp2 cell responses in the absence (left) or presence (right) of 100 μm Mianserin, an antagonist of *Drosophila* 5-HT2 receptor subtypes (Blenau et al., 2017; Colas et al., 1995; Tierney, 2018; Vleugels et al., 2015). (I) Mean ΔF/F_0_ traces (black lines) ± SEM (gray shading). Blue bars indicate proboscis sucrose stimulation. (J) Scatter plots of maximum ΔF/F_0_ changes, with bar plot overlay, mean ± SEM. n = 9 flies; paired t-test, ** p < 0.01. (K-N) Dilp2 cells respond to optogenetic excitation of sugar-SEL PNs. (K) Experiment schematic. CsChrimson (CsChR) was expressed in sugar-SEL PNs while GCaMP6s (GC6s) was independently expressed in Dilp2 cells. (L) Representative GCaMP6s ΔF/F_0_ images of Dilp2 cell responses in the absence (left) or presence (right) of 100 μm Mianserin. (M) Mean ΔF/F_0_ traces (black lines) ± SEM (gray shading). Red bars indicate red light stimulations. (N) Scatter plots of maximum ΔF/F_0_ changes, with bar plot overlay, mean ± SEM. n = 7 flies; paired t-test, *** p < 0.001. (O and P) Co-labelling of sugar-SEL LNs (*SEZ-569>UAS-CD8-RFP*, magenta) and Dilp2 insulin-producing cells (*Dilp2-LexA>LexAop-CD8-GFP*, green). Anti-Brp staining shows brain neuropil (blue). Yellow asterisks in (O) mark other cells labelled by *SEZ-569*. Scale bar, 50 μm. (Q and R) Dilp2 cells do not respond to optogenetic excitation of sugar-SEL LNs. (Q) Mean ΔF/F_0_ traces (black lines) ± SEM (gray shading) of Dilp2 cells in experimental (*SEZ-569>CsChrimson*) and control flies (*UAS-CsChrimson*). Red bars indicate red light stimulations. (R) Scatter plots of maximum ΔF/F_0_ changes of the indicated genotypes, with bar plot overlay, mean ± SEM. n = 8 *SEZ-569* flies, n = 7 *UAS* flies; Mann Whitney test, ns, not significant. See Figure S2 for additional characterizations of Dilp2 cells.

To investigate whether the insulin-producing cells express 5-HT receptors and therefore might directly detect sugar-SEL 5-HT release, we examined 5-HT receptor expression in the PI using knock-in *Gal4* lines (Gnerer et al., 2015). Of the five 5-HT receptors in *Drosophila*, only 5-HT2A is expressed in a subset of insulin-producing cells, albeit at low levels (Figures S2B-S2F’’ and 3D-3F). 5-HT2A is an excitatory 5-HT receptor (Vleugels et al., 2015), suggesting that insulin-producing cells are activated by sugar-SEL PNs. Consistent with this hypothesis, we found that insulin-producing cells responded to sugar taste detection (Figures 3G-3I), similar to sugar-SELs. The response persisted in flies with the esophagus severed (Figures S2G-S2I), indicating that the response was independent of sugar ingestion. This is reminiscent of the cephalic phase insulin response in mammals, where insulin is secreted in response to the sensory detection of food well before blood glucose levels change (Power and Schulkin, 2008; Teff, 2000; Zafra et al., 2006). In contrast, Diuretic hormone 44 (Dh44) expressing neurons in the PI, which sense circulating glucose (Dus et al., 2015), were not acutely excited by sugar taste detection (Figures S2J-S2L).

To test the hypothesis that sugar-SELs relay sugar taste information to the insulin-producing cells through the activation of 5-HT2A, we applied mianserin, an antagonist of *Drosophila* 5-HT2 receptor subtypes (Blenau et al., 2017; Colas et al., 1995; Tierney, 2018; Vleugels et al., 2015), to the exposed fly brain. The presence of mianserin diminished taste-evoked calcium responses in insulin-producing cells (Figures 3H-3J). To directly test the hypothesis that sugar-SELs excite the insulin-producing cells, we activated sugar-SEL PNs using CsChrimson, while simultaneously monitoring the response in insulin-producing cells (Figure 3K). Optogenetic activation of sugar-SEL PNs consistently excited the insulin-producing cells, demonstrating a functional excitatory connection (Figures 3L-3N). This excitation was largely inhibited in the presence of mianserin (Figures 3L-3N), consistent with a requirement for 5-HT2A. In contrast, sugar-SEL LNs have little anatomical overlap with the insulin-producing cells (Figures 3O and 3P), and optogenetic activation of sugar-SEL LNs did not consistently excite insulin-producing cells (Figures 3Q and 3R). Taken together, these results indicate that sugar-SEL PNs relay sugar taste detection to insulin-producing cells, likely to promote insulin release in anticipation of sugar consumption.

### Sugar-SELs Limit Sugar Consumption

The findings that sugar-SELs respond to sugar taste detection on the proboscis but not on the legs (Figures 2N-2P) and that they activate insulin-producing cells (Figures 3K-3N) suggest that they play a role specifically related to feeding. To test the hypothesis that sugar-SELs regulate feeding, we measured sugar intake in fasted flies, using a temporal consumption assay (Pool et al., 2014). Transient excitation of the sugar-SELs using the heat-activated cation channel dTRPA1 (Hamada et al., 2008) immediately before and during the consumption assay decreased sugar consumption (Figures 4A and 4B). In contrast, a transient inhibition of synaptic transmission in sugar-SELs by overexpressing a dominant-negative, temperature-sensitive Dynamin (*Shi^ts^*) (Kitamoto, 2001), increased sugar consumption (Figures 4C and 4D). At permissive temperatures where dTRPA1 and Shi^ts^ were not active, sugar intake remained unchanged (Figure S3). Importantly, blocking 5-HT synthesis in sugar-SELs, by specific expression of *Tryptophan hydroxylase* (*Trh*) RNAi in these cells, also increased sugar consumption (Figure 4E). These findings demonstrate that the 5-HT output from sugar-SELs limits sugar intake.

**Figure 4.**
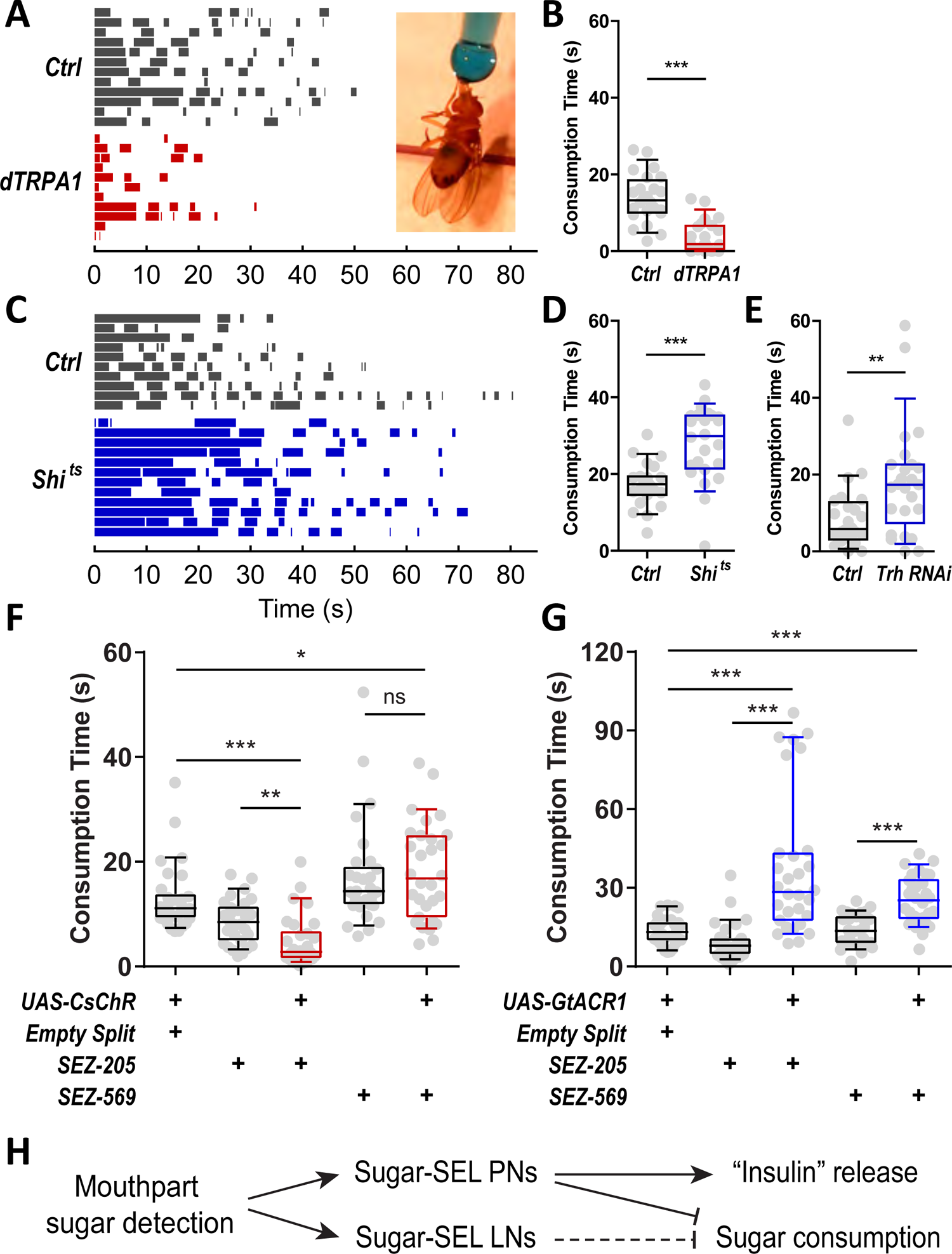
Sugar-SELs Limit Sugar Intake. (A and B) Representative feeding bouts (A) and total consumption time (B) of control flies (*Ctrl*) and flies upon transient excitation of sugar-SELs (*dTRPA1*). Inset (A) shows a fly drinking from a sucrose droplet (blue). n = 24 flies/genotype; Mann Whitney test, *** p < 0.001. (C and D) Representative feeding bouts (C) and total consumption time (D) of control flies (*Ctrl*) and flies upon transient silencing of sugar-SELs (*Shi^ts^*). n = 23 *Ctrl* flies, n = 24 *Shi^ts^* flies; Mann Whitney test, *** p < 0.001. (E) Total consumption time of control flies (*Ctrl*) and flies with *Trh* RNAi in sugar-SELs (*Trh RNAi*). n = 24 *Ctrl* flies, n = 25 *Trh RNAi* flies; Mann Whitney test, ** p < 0.01. (F) Total consumption time of control flies and flies with sugar-SEL PNs (*SEZ-205*) or sugar-SEL LNs (*SEZ-569*) transiently excited using CsChrimson. n = 30 flies/genotype; Kruskal-Wallis ANOVA followed by Dunn’s multiple comparisons tests, * p < 0.05; ** p < 0.01; *** p < 0.001; ns, not significant. (G) Total consumption time of control flies and flies with sugar-SEL PNs (*SEZ-205*) or sugar-SEL LNs (*SEZ-569*) transiently silenced using GtACR1. n = 30 flies/genotype; Kruskal-Wallis ANOVA followed by Dunn’s multiple comparisons tests, *** p < 0.001. (H) A summary model of the functions of sugar-SEL PN and LNs. Horizontal bars in (A) and (C) show feeding bouts of individual flies (one line per fly). For all box plots, whiskers = 10^th^ to 90^th^ percentile, box = 25^th^ to 75^th^ percentile, and line in box = median. Dots indicate individual data. See Figure S3 for dTRPA1 and Shi^ts^ control experiments.

Given that sugar-SEL PNs and LNs have different anatomy and functional connectivity to insulin-producing cells (Figures S1A-S1G and 3K-3R), we examined whether they differentially regulate sugar consumption. Specific optogenetic excitation of the sugar-SEL PNs decreased sugar consumption (Figure 4F, *SEZ-205*), whereas specific excitation of the sugar-SEL LNs had no measurable effect (Figure 4F, *SEZ-569*). However, hyperpolarizing either sugar-SEL PNs or LNs using the light-gated anion channel GtACR1 (Mohammad et al., 2017) increased sugar consumption (Figure 4G), similar to the effect of silencing all sugar-SELs (Figures 4C and 4D). These results indicate that sugar-SEL PNs and LNs cooperate to limit sugar consumption (Figure 4H). Taken together, our studies demonstrate that activation of sugar-SELs by sugar gustatory detection increases the activity of insulin-producing cells and decreases consumption, providing insight into the neural circuit that regulates internal nutrient availability and consumption in anticipation of sugar intake.

### Bitter-SELs Respond to Bitter Detection on Proboscis and Legs

To elucidate the function of all SEZ 5-HT neurons that respond to taste detection, we next examined the role of bitter-SELs in gustatory processing and feeding regulation. The distinct morphology of bitter-SELs (Figures 1A and 1B, lower panels) allowed us to visually identify a *Gal4* driver, *VT46202-Gal4*, that labels them. *VT46202-Gal4* has sparse expression in the central brain, labeling only the bitter-SELs in the SEZ (Figure S4A and S4A’). The genetic intersection of this line and *Dfd-LexA* restricted *Gal4* expression to two pairs of 5-HT neurons, the bitter-SELs (Figures 5A and 5B; Video S1). In addition to arborizing in the SEZ, bitter-SELs send descending processes that arborize on the dorsal surface of the ventral nerve cord (Figures 5A, 5B, S4C, and S4C’). Based on this morphology, we identified bitter-SELs as DNg28 descending neurons (DN) in the DN *split-Gal4* collection (Namiki et al., 2018) (Figures S4B and S4B’). The dendrites of bitter-SELs are largely restricted to the SEZ, consistent with receiving taste input, and the descending projections are axonal (Figure 5C).

**Figure 5.**
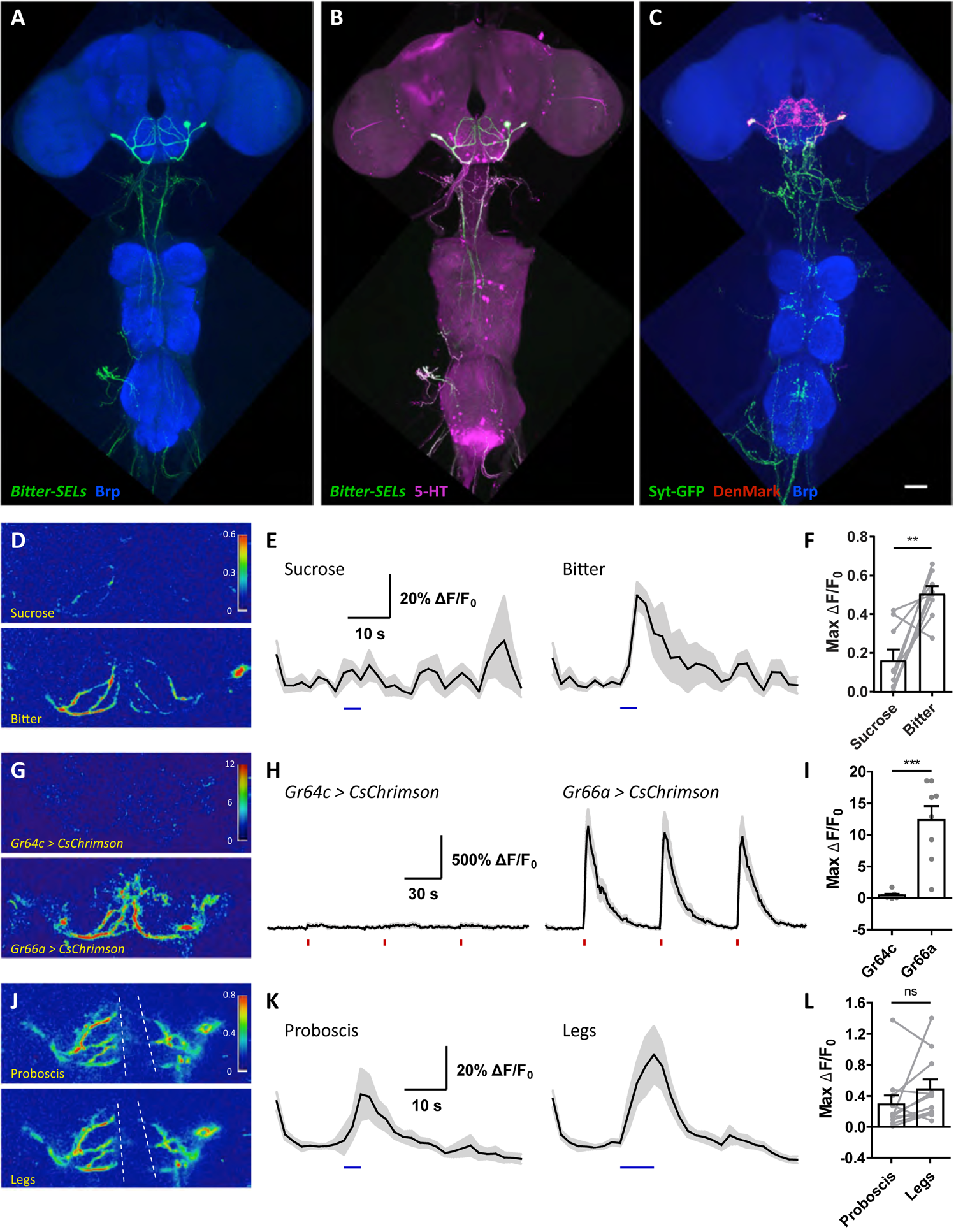
Bitter-SELs Respond to Proboscis and Leg Bitter Detection. (A and B) Bitter-SELs are labelled by the genetic intersection between *VT46202-Gal4* and *Dfd-LexA* (green). Anti-Brp staining (A) shows brain and VNC neuropil (blue). Anti-5-HT staining (B) shows 5-HT neurons (magenta). (C) Dendritic (DenMark, red) and axonal (Syt-GFP, green) compartments of bitter-SELs. Anti-Brp staining shows brain and VNC neuropil (blue). Scale bar, 50 μm. (D-F) Bitter-SELs preferentially respond to proboscis bitter taste detection. (G-I) Bitter-SELs respond to CsChrimson-mediated optogenetic excitation of bitter (*Gr66a^+^*) but not sugar (*Gr64c^+^*) GRNs. (J-L) Bitter-SELs respond to bitter detection on the proboscis and the legs. Images in (D), (G), and (J) are representative ΔF/F_0_ images of GCaMP6s responses. White dotted lines in (J) indicate the esophagus. Plots in (E), (H), and (K) show mean ΔF/F_0_ traces (black lines) ± SEM (gray shading). Blue bars (E) indicate proboscis sucrose (left) or bitter (right) stimulation. Red bars (H) indicate red light stimulations. Blue bars (K) indicate bitter stimulation on the proboscis (left) or legs (right). (F), (I), and (L) are scatter plots of maximum ΔF/F_0_ changes, with bar plot overlay, mean ± SEM. (F): n = 9 flies; paired t-test, ** p < 0.01. (I): n = 7 *Gr64c* flies, n = 8 *Gr66a* flies; Mann Whitney test, *** p < 0.001. (L): n = 11 flies; paired t-test, ns, not significant. See Video S1 for bitter-SEL anatomy. See Figure S4 for additional characterization of bitter-SELs.

Using specific drivers to express GCaMP6s in bitter-SELs, we confirmed that they responded to bitter taste detection on the proboscis (Figures 5D-5F), consistent with the *Trh-Gal4* studies (Figures 1A and 1B). Moreover, bitter-SELs were consistently activated by optogenetic excitation of bitter (*Gr66a^+^*) but not sugar (*Gr64c^+^*) GRNs (Figures 5G-5I). In contrast to the specific response of sugar-SELs to proboscis sugar detection (Figures 2N-2P), bitter-SELs responded to bitter detection on both the proboscis and the legs (Figures 5J-5L). These findings suggest that bitter-SELs may play a general role in bitter taste detection rather than a specific role in regulating consumption. Indeed, acute excitation of bitter-SELs had no measurable effect on food intake (Figures S4D and S4E).

### Bitter-SELs Excite 5-HT7-Expressing Enteric Neurons

In addition to the arborizations in the central nervous system, bitter-SELs have processes associated with the digestive system. Bitter-SELs send axons through the labial nerves (Figures 6B-6D and S5B) and along the esophagus (Figure S5B) to the hypocerebral ganglion (Figures 6A-6D and S5A). Interestingly, bitter-SEL projections account for all of the 5-HT processes that innervate the hypocerebral ganglion (Figures 6E-6E’’). The hypocerebral ganglion is an enteric ganglion at the junction of the esophagus, the crop (food storage), and the proventriculus/anterior midgut (food grinding and digestion) (Figure 6A) (Lemaitre and Miguel-Aliaga, 2013; Miguel-Aliaga et al., 2018). This junction region is a key site for food passage regulation as it determines whether food enters the crop for temporary storage or the midgut for digestion. Therefore, the anatomy of bitter-SEL projections suggests the hypothesis that they participate in the regulation of digestion.

**Figure 6.**
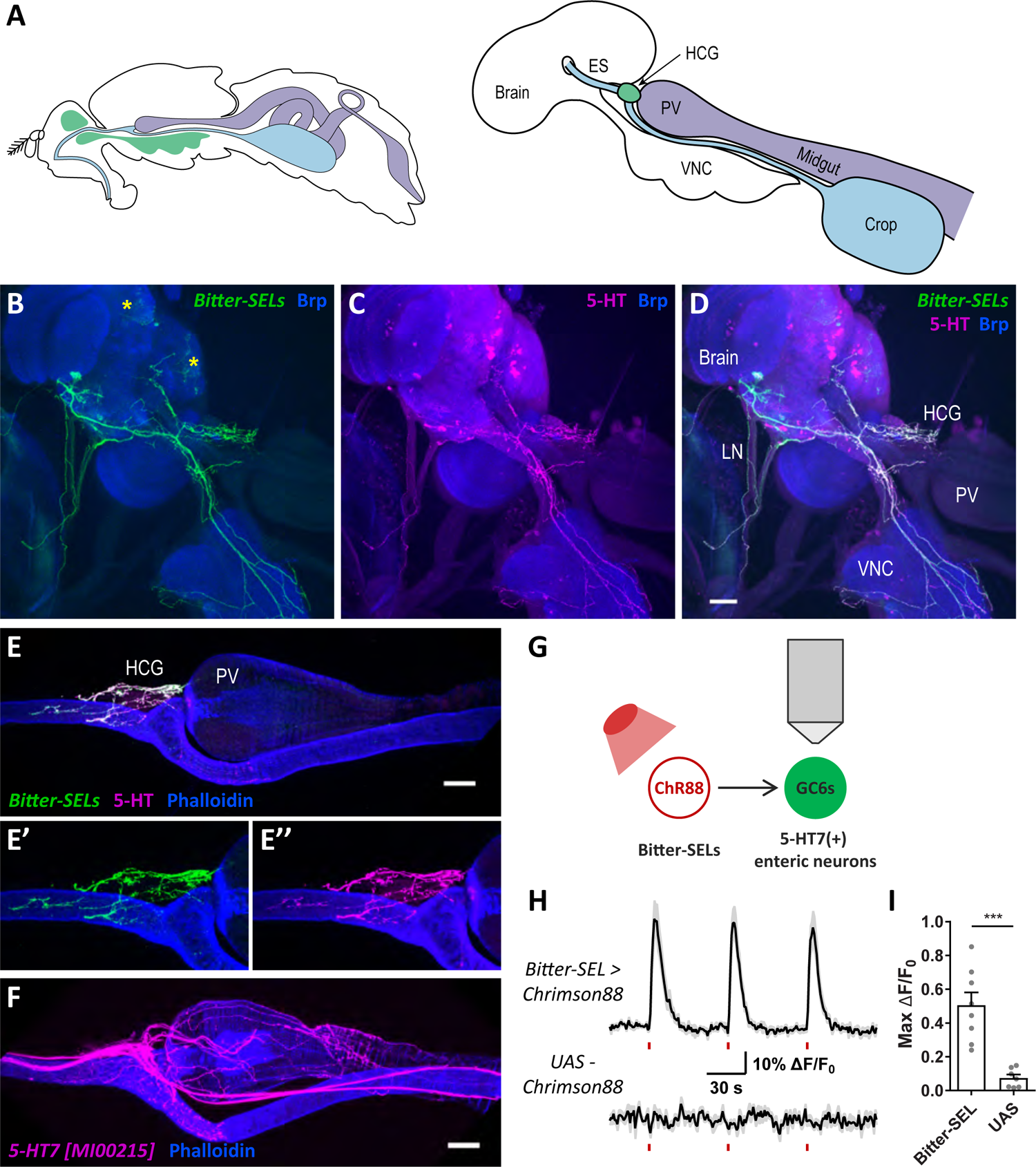
Bitter-SELs Activate 5-HT7 Expressing Enteric Neurons. (A) Left: A diagram of the central nervous system (green), esophagus and crop (blue), and the rest of the digestive tract (purple). Right: A diagram of the relative position of the central nervous system and digestive tract as shown in (B-D). VNC, ventral nerve cord; ES, esophagus; PV, proventriculus; HCG: hypocerebral ganglion. (B-D) Bitter-SEL processes are labelled by *DN052>UAS-CD8-GFP* and stained for GFP (green). Yellow asterisks (B) mark non-specific cells labelled by *DN052*. Anti-5-HT staining shows 5-HT neurons and processes (magenta). Anti-Brp staining shows brain and VNC neuropil (blue). LN, labial nerves. Scale bar, 50 μm. (E-E’’) A close-up view of bitter-SEL processes (green) in the hypocerebral ganglion (HCG) in a different fly. Anti-5-HT staining (magenta); Phalloidin stains the digestive tract (blue). Scale bar, 50 μm. (F) 5-HT7 expressing enteric neurons are labelled by *5-HT7[MI00215]-Gal4>UAS-CD8-RFP* and stained for RFP (magenta). Phalloidin stains the digestive tract (blue). Scale bar, 50 μm. (G-I) Optogenetic excitation of bitter-SELs activates 5-HT7 expressing enteric neurons. (G) Experiment schematic. Chrimson88 (ChR88) was expressed in bitter-SELs while GCaMP6s (GC6s) was independently expressed in 5-HT7(+) enteric neurons. (H) Mean ΔF/F_0_ traces (black lines) ± SEM (gray shading) of 5-HT7(+) enteric neurons in experimental flies (*Bitter-SEL>UAS-Chrimson88*) and control flies (*UAS-Chrimson88*). Red bars indicate red light stimulations. (I) Scatter plots of maximum ΔF/F_0_ changes of the indicated genotypes, with bar plot overlay, mean ± SEM. n = 8 *Bitter-SEL* flies, n = 7 *UAS* flies; Mann Whitney test, *** p < 0.001. See Figure S5 for additional analysis on bitter-SELs, AKH cells, and 5-HT receptor expression in HCG.

To examine the possibility that bitter-SELs synapse onto enteric neurons in the proventricular region and regulate digestion, we first examined the expression of 5-HT receptors in this region using knock-in *Gal4* lines (Gnerer et al., 2015). Of the five 5-HT receptors, the excitatory 5-HT receptor 5-HT7 is expressed most abundantly in the proventricular region (Figures S5C-S5G’’), in neuronal processes innervating the proventriculus/anterior midgut and the crop (Figures 6F and S5G’). These 5-HT7(+) enteric processes are thus well positioned to receive 5-HT input from the bitter-SELs (Figures 6E-6F). Consistent with this, we found that optogenetic excitation of bitter-SELs induced calcium increases in the 5-HT7(+) enteric processes, demonstrating a functional excitatory connection (Figures 6G-6I). In contrast, nearby Adipokinetic hormone (AKH) cells did not express detectable levels of 5-HT receptors (Figures S5C-S5G’’) and were not activated by acute optogenetic excitation of bitter-SELs (Figures S5H-S5J). In summary, bitter-SELs innervate the hypocerebral ganglion and excite 5-HT7(+) enteric neurons in the proventricular region, suggesting that they regulate enteric physiology and function.

### Bitter-SELs and 5-HT7-Expressing Enteric Neurons Promote Crop Contractions

5-HT modulates gastrointestinal motility in mammals and insects (Tecott, 2007). In insects including *Drosophila*, 5-HT promotes crop (food storage organ) contractions (e.g., (Calkins et al., 2017; French et al., 2014; Liscia et al., 2012; Solari et al., 2017)); however, the source of 5-HT and the underlying 5-HT receptors are unknown. Our finding that bitter-SELs project to the hypocerebral ganglion and excite 5-HT7(+) enteric neurons (Figure 6) suggests that bitter-SELs may supply 5-HT to the digestive tract to modulate crop contractions via 5-HT7(+) enteric neurons. To test this, we developed a preparation to observe crop contractions in live flies that is compatible with simultaneous optogenetic stimulation (Figure 7A; STAR Methods).

**Figure 7.**
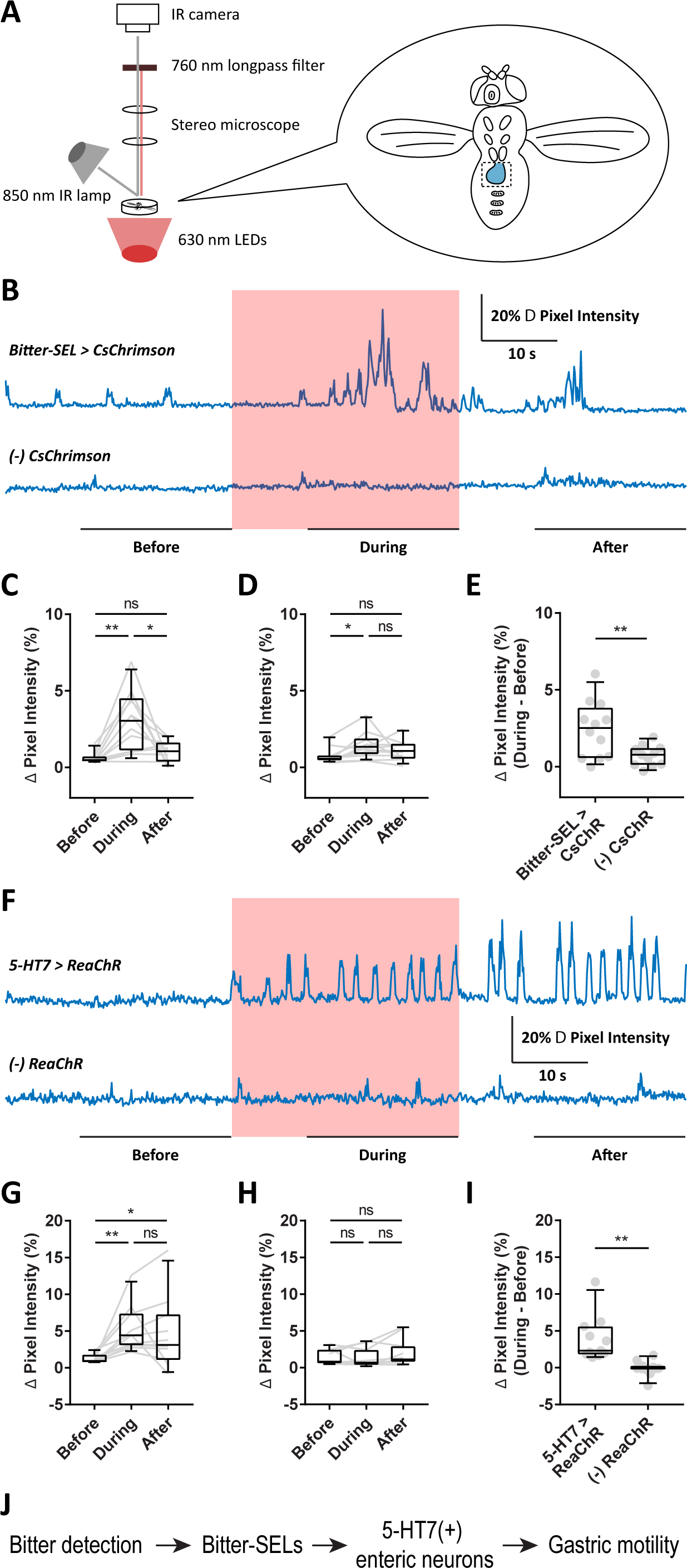
Optogenetic Excitation of Bitter-SELs and 5-HT7 Expressing Enteric Neurons Promote Crop Contractions. (A) Schematic of the assay to video-record crop contractions in live flies in conjunction with simultaneous optogenetic stimulation. (B) Representative traces of crop contractions in an experimental fly with CsChrimson expression in bitter-SELs (*Bitter-SEL>CsChrimson*) and a control fly (*(-)CsChrimson*). The red-shaded area indicates 630 nm light stimulation. (C and D) Average crop contractions before, during, and after 630 nm light stimulation (time windows as illustrated in (B)) for experimental *Bitter-SEL>CsChrimson* flies (n = 12) (C) and control *(-)CsChrimson* flies (n = 11) (D). Friedman test with Dunn’s multiple comparisons test, * p < 0.05; ** p < 0.01; ns, not significant. (E) The net increase of crop contractions (During - Before) for experimental *Bitter-SEL>CsChrimson* flies (n = 12) and control *(-)CsChrimson* flies (n = 11). Unpaired t test with Welch’s correction, ** p < 0.01. (F) Representative traces of crop contractions in an experimental fly with ReaChR expression in 5-HT7 neurons (*5-HT7>ReaChR*) and a control fly (*(-)ReaChR*). The red-shaded area indicates 630 nm light stimulation. (G and H) Average crop contractions before, during, and after 630 nm light stimulation (time windows as illustrated in (F)) for experimental *5-HT7>ReaChR* flies (n = 11) (G) and control *(-)ReaChR* flies (n = 11) (H). Friedman test with Dunn’s multiple comparisons test, * p < 0.05; ** p < 0.01; ns, not significant. (I) The net increase of crop contractions (During - Before) for experimental *5-HT7>ReaChR* flies (n = 11) and control *(-)ReaChR* flies (n = 11). Unpaired t test with Welch’s correction, ** p < 0.01. (J) Summary model for bitter-SEL function. For all box plots, whiskers = 10^th^ to 90^th^ percentile, box = 25^th^ to 75^th^ percentile, and line in box = median. See Video S2 for an example of crop contractions induced by bitter-SEL excitation.

We found that optogenetic excitation of bitter-SELs using CsChrimson acutely increased crop contractions during the stimulation period (Figures 7B-7E). Similarly, when 5-HT7(+) neurons were activated using the light-gated cation channel ReaChR (Inagaki et al., 2014; Lin et al., 2013) (because CsChrimson expression induced lethality), crop contractions increased relative to controls and were sustained for tens of seconds post-stimulation (Figures 7F-7I). These studies show that both bitter-SELs and downstream enteric targets promote crop contractions. As regurgitation was not observed when bitter-SELs were excited during the consumption assay (Figures S4D and S4E), these results suggest that bitter-SEL excitation promotes the movement of food from the crop into the midgut for digestion. Taken together, our findings suggest the model that under conditions of poor food quality (i.e., in the presence of bitter and thus potentially harmful substances), bitter-SELs activate 5-HT7(+) enteric neurons to promote crop contractions, thereby utilizing crop food reserves for energy replenishment in anticipation of potential food shortages (Figure 7J). Thus, our studies show that both sugar-SELs and bitter-SELs transform taste detection into anticipatory regulation of internal nutrient availability.

## Discussion

In this study, we identified two classes of 5-HT neurons in the fly gustatory center that play distinct roles in gustatory processing and function independently to promote nutrient homeostasis. One class, the sugar-SELs, responds to sugar gustatory detection, promotes insulin-producing cell activity and reduces feeding drive, suggesting that they prevent overconsumption in nutrient rich environments. The other class, the bitter-SELs, responds to sensory detection of bitter compounds and promotes crop contractions, likely to utilize stored food in nutrient poor environments. Thus, 5-HT neurons are essential circuit elements that coordinate endocrine and digestive function in anticipation of altered food intake.

### 5-HT Has Multifaceted Roles in Feeding Regulation and Nutrient Homeostasis

5-HT profoundly modulates appetite and feeding across animal species (Tecott, 2007). In humans and rodents, the global effect of brain 5-HT signaling is the suppression of food intake; however, the involvement of multiple brain regions (including the hypothalamus, the nucleus of the solitary tract, and the parabrachial nuclei) and multiple 5-HT receptors (e.g., 5-HT1B, 5-HT2C, 5-HT6) underscores the complex nature of 5-HT modulation of appetite and feeding (Donovan and Tecott, 2013; Lam et al., 2010; Marston et al., 2011; Tecott, 2007; Wyler et al., 2017). Moreover, the complexity of the mammalian 5-HT network provides challenges in elucidating the specific 5-HT circuits that critically regulate feeding. Studies in invertebrate models have likewise demonstrated multiple, sometimes opposite, roles for 5-HT neurons in regulating feeding (Tierney, 2020). For instance, in *Drosophila* adults, activating all 5-HT neurons suppresses food intake whereas activating a smaller yet diverse subset promotes food intake, suggesting heterogeneity in 5-HT regulation of feeding (Albin et al., 2015; Pooryasin and Fiala, 2015). With the exception of a few cases where specific 5-HT neurons that influence feeding have been identified – for example, the 5-HT neurons that promote pharyngeal pumping rate to enhance food ingestion in *C. elegans* and *Drosophila* larvae (Ishita et al., 2020; Schoofs et al., 2018; Tierney, 2020) – the diversity of 5-HT neurons that contributes to feeding regulation and their influence on nutrient stores remains largely uncharacterized.

Here, we identify multiple classes of 5-HT neurons that are activated by gustatory detection and signal to the endocrine and digestive systems to influence nutrient availability. Sugar-SEL PNs activate insulin-producing cells and decrease consumption (Figures 3 and 4), with feeding regulation occurring either via insulin-producing cells or an independent pathway. Silencing sugar-SEL LNs also increases consumption (Figure 4G), arguing that sugar taste detection activates local and projecting 5-HT pathways that limit consumption. In contrast, bitter-SELs have no measurable effect on food consumption (Figures S4D and S4E) but instead activate enteric neurons that elicit crop contractions (Figures 6 and 7), suggesting that they influence nutrient availability by mobilizing stored nutrients. Thus, our studies reveal 5-HT neurons that have different projection patterns, relay sugar and bitter gustatory information to different downstream targets, and regulate internal nutrient availability by distinct mechanisms. These studies provide insight into the multifaceted roles of 5-HT neurons in gustatory processing, feeding regulation, and nutrient homeostasis, highlighting the importance of understanding the myriad functions of 5-HT at the neural circuit level.

### Sugar Taste Detection Preemptively Regulates Insulin Release and Feeding Drive

Food-derived sensory cues of food are used as anticipatory signals to regulate endocrine function and feeding drive. Studies in mammalian models have revealed that sensory detection of food triggers preparatory (cephalic phase) insulin release (Power and Schulkin, 2008; Teff, 2000; Zafra et al., 2006) and rapidly inhibits the activity of Agouti-Related Peptide (AgRP) hunger neurons (Betley et al., 2015; Chen and Knight, 2016; Chen et al., 2015; Mandelblat-Cerf et al., 2015). The neural circuits that underlie these preparatory mechanisms, however, are not fully understood.

Our findings suggest that insulin release in anticipation of food consumption is a process shared in flies and mammals, perhaps indicating an effective strategy to promote rapid nutrient storage during feeding and to prevent overconsumption. Using *in vivo* calcium imaging, we found that fly insulin-producing cells are rapidly excited by sugar gustatory detection independent of consumption (Figures 3G-3J and S2G-S2I). Moreover, we identified the sugar-SELs as a specific neural pathway that mediates the preparatory insulin response. The 5-HT output from these neurons exerts an inhibitory effect on sugar intake, suggesting that it anticipates nutrient influx and serves as a feedback mechanism to limit consumption (Figure 4). Thus, we have identified a specific class of 5-HT neurons that mediates the preparatory insulin response as well as the reduction of feeding drive in response to sugar gustatory cues, shedding light on the neural circuit mechanisms for anticipatory responses to food detection.

### Bitter Gustatory Detection May Be Used to Predict Food Scarcity

A surprising finding from our study is that bitter-SELs, which respond to bitter gustatory detection, promote contractions of the crop food storage organ. While there is evidence that intestinal bitter detection modulates gastrointestinal physiology (Sarnelli et al., 2019; Xie et al., 2018), the regulation of gastrointestinal function by bitter gustatory detection is less examined. We reason that because bitter compounds are often toxic, encounters with bitter compounds may indicate poor feeding conditions and potential food shortage. Under such conditions, bitter-SELs may function to promote crop motility to utilize food reserves in anticipation of limited food intake. We therefore propose that bitter gustatory compounds may have an unappreciated role in predicting food scarcity and stimulating digestion as a preparatory response.

In addition to the hypocerebral ganglion, bitter-SELs project to diverse targets (Figures 6B-6D, S5A, and S5B), suggesting that they may carry out other functions besides enteric modulation. For example, bitter-SELs broadly arborize on the dorsal surface of the ventral nerve cord (Figures S4C, S4C’, and S5A) (Namiki et al., 2018). Thus, bitter-SELs may also set the 5-HT tone in the ventral nerve cord or secrete 5-HT into the hemolymph to modulate target tissues in a paracrine or endocrine fashion.

### Neuromodulatory Circuits as Candidates for Mediating Preparatory Responses

We find that 5-HT neurons are critical nodes in the circuits that transform acute gustatory detection into longer-term changes in endocrine and digestive function. Although the timescale of activation of sugar-SELs and bitter-SELs and the dynamics of 5-HT release requires further investigation, 5-HT receptors are metabotropic receptors ideally suited for transforming transient neural signals into more sustained cellular responses. In this regard, neuromodulatory circuits are prime candidates for eliciting preparatory responses that require the transformation of neural signals across time scales. Our work thus sheds light on neural circuit mechanisms that translate external sensory cues into preparatory physiological responses and suggests that neuromodulators such as 5-HT may contribute to anticipatory mechanisms in other animals.

## Acknowledgments

We thank Christoph Scheper for identifying *VT46202-Gal4* and the *split-Gal4* line for the bitter-SELs, Gabriella R. Sterne for generating and identifying *split-Gal4* lines for sugar-SEL PNs and LNs, Brendan C. Mullaney for generating *AKH-LexA* and members of the Scott lab for helpful discussion and feedback. Two-photon imaging experiments were conducted at the CRL Molecular Imaging Center, supported by NSF DBI-1041078 and the Helen Wills Neuroscience Institute. We thank Julie H. Simpson, Hubert Amrein, David J. Anderson, Adam Claridge-Chang, Herman A. Dierick, Zhefeng Gong, Anthony Cammarato, Barry J. Dickson, Paul A. Garrity, Toshihiro Kitamoto, the Bloomington *Drosophila* Stock Center, and the Vienna *Drosophila* Resource Center for fly stocks. This work was supported by a Jane Coffin Childs Fellowship to Z.Y. and an NIH grant R01GM128209 to K.S.

## Author Contributions

Z.Y. conceived and performed experiments and analyzed the data under guidance from K.S. Z.Y. and K.S. wrote the manuscript.

---

**Figure S1.**
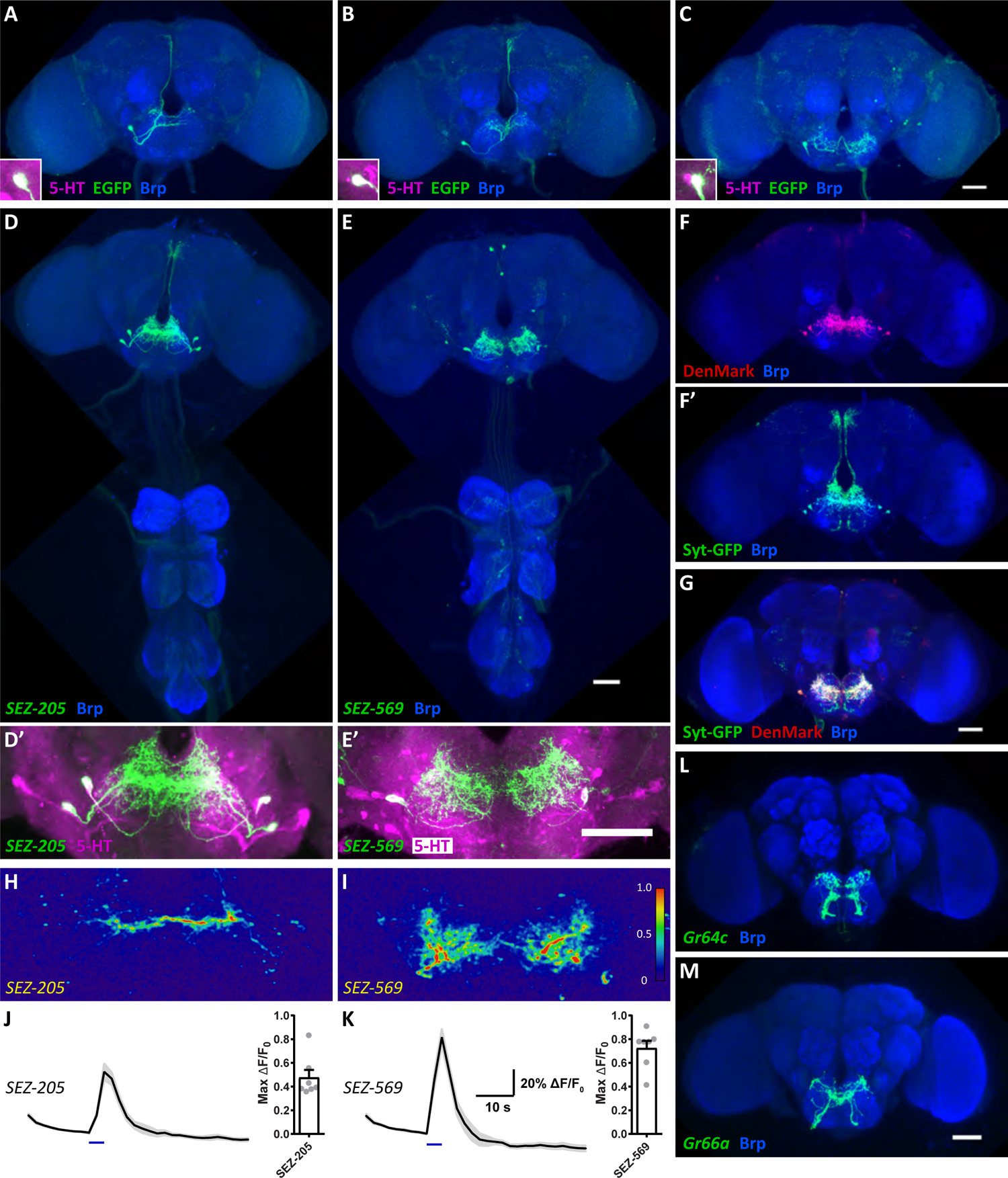
Subtypes of Sugar-SELs, Related to Figure 2. (A-C) Representative single-cell morphology of sugar-SELs showing an ipsilateral PN (A), a contralateral PN (B), and an LN (C) (green). Insets show co-staining against 5-HT (magenta). Anti-Brp staining shows brain neuropil (blue). Scale bar, 50 μm. (D and D’) The *SEZ-205 split-Gal4* specifically labels two pairs of sugar-SEL PNs (green). Anti-5-HT staining (magenta) and anti-Brp staining (blue). (E and E’) The *SEZ-569 split-Gal4* labels sugar-SEL LNs and other neurons (green). Anti-5-HT staining (magenta) and anti-Brp staining (blue). Scale bars, 50 μm. (F and F’) Dendritic (DenMark) and axonal (Syt-GFP) compartments of sugar-SEL PNs. Anti-Brp staining shows brain neuropil (blue). (G) Dendritic (DenMark) and axonal (Syt-GFP) compartments of sugar-SEL LNs. Anti-Brp staining shows brain neuropil (blue). Scale bar, 50 μm. (H and I) Representative GCaMP6s ΔF/F_0_ images of sugar-SEL PNs (H) and sugar-SEL LNs (I) upon proboscis sucrose detection. (J and K) Left: mean ΔF/F_0_ traces (black lines) ± SEM (gray shading) for sugar-SEL PNs (J) and sugar-SEL LNs (K) upon proboscis sucrose detection (blue bars). Right: scatter plots of maximum ΔF/F_0_ changes for individual flies (n = 7 flies/genotype), with bar plot overlay, mean ± SEM. (L and M) Expression of CsChrimson in sugar (L) and bitter (M) GRNs (green) for optogenetic experiments shown in Figures 2K-2M and 5G-5I. Anti-Brp staining shows brain neuropil (blue). Scale bar, 50 μm.

**Figure S2.**
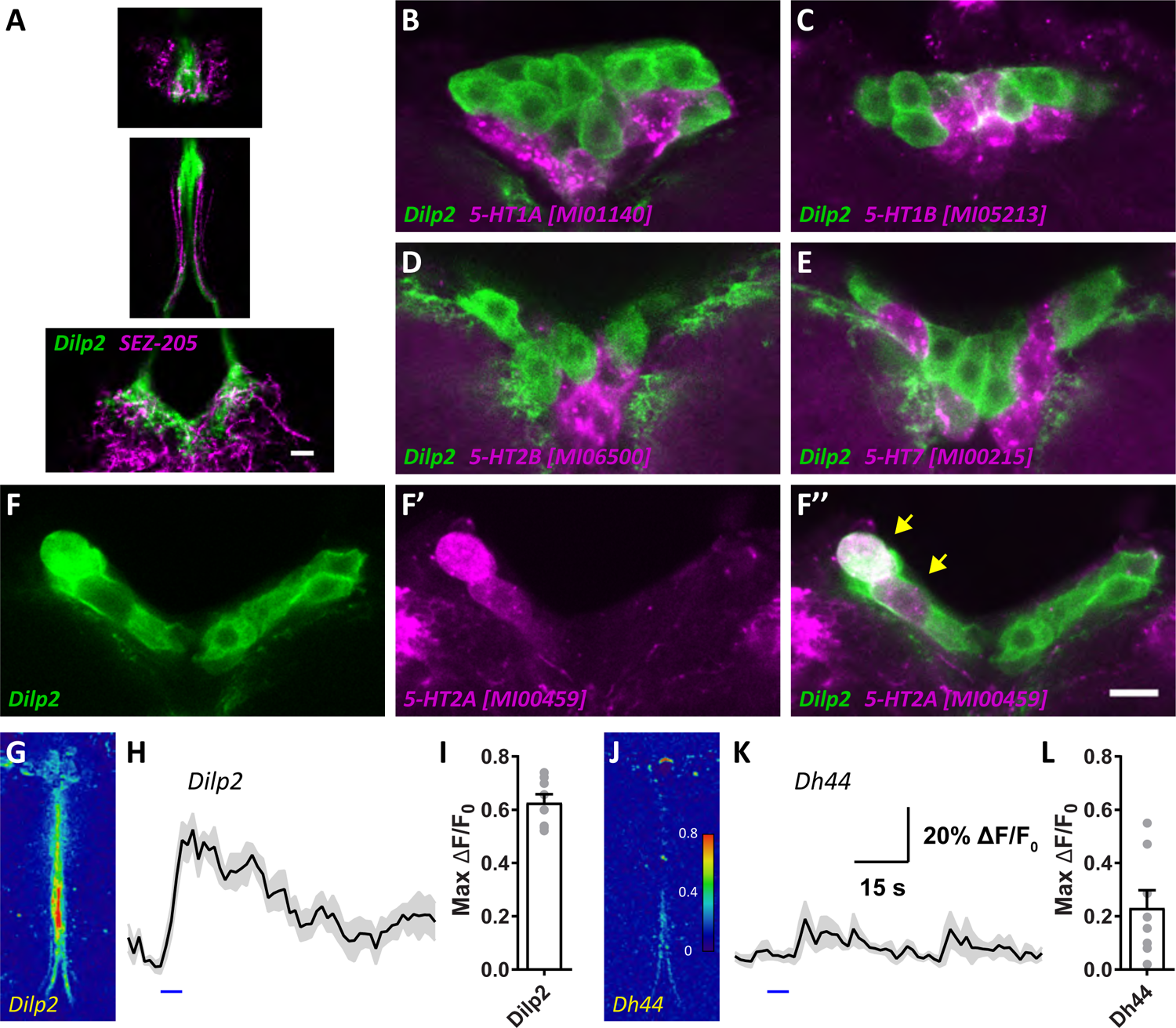
Additional Analysis of Insulin-Producing Cells, Related to Figure 3. (A) Co-labelling of Dilp2 insulin-producing cells (*Dilp2-LexA>LexAop-CD8-GFP*, green) and sugar-SEL PNs (*SEZ-205>UAS-CD8-RFP*, magenta) in single confocal slices (1.3 μm thick) near the PI (top), along the median bundle (middle), and in the SEZ (bottom). These images are from the same brain shown in Figures 3A-3C. Scale bar, 10 μm. (B-F’’) Representative images showing co-labelling of Dilp2 insulin-producing cells (*Dilp2-LexA>LexAop-CD8-GFP*, green) and the indicated 5-HT receptors (*5-HT receptor Gal4>UAS-CD8-RFP*, magenta) in single confocal slices (1.3 μm thick). Scale bar, 10 μm. (G-I) Dilp2 insulin-producing cells respond to proboscis sucrose detection in flies with the esophagus severed. (J-L) Dh44 neurons do not consistently respond to proboscis sucrose detection. Images in (G) and (J) are representative GCaMP6s ΔF/F_0_ images of Dilp2 cells (G) and Dh44 neurons (J) in response to proboscis sucrose detection. Plots in (H) and (K) show mean ΔF/F_0_ traces (black lines) ± SEM (gray shading). Blue bars in (H) and (K) indicate proboscis sucrose stimulation. (I) and (L) are scatter plots of maximum ΔF/F_0_ changes for individual flies (n = 8 flies/genotype), with bar plot overlay, mean ± SEM.

**Figure S3.**
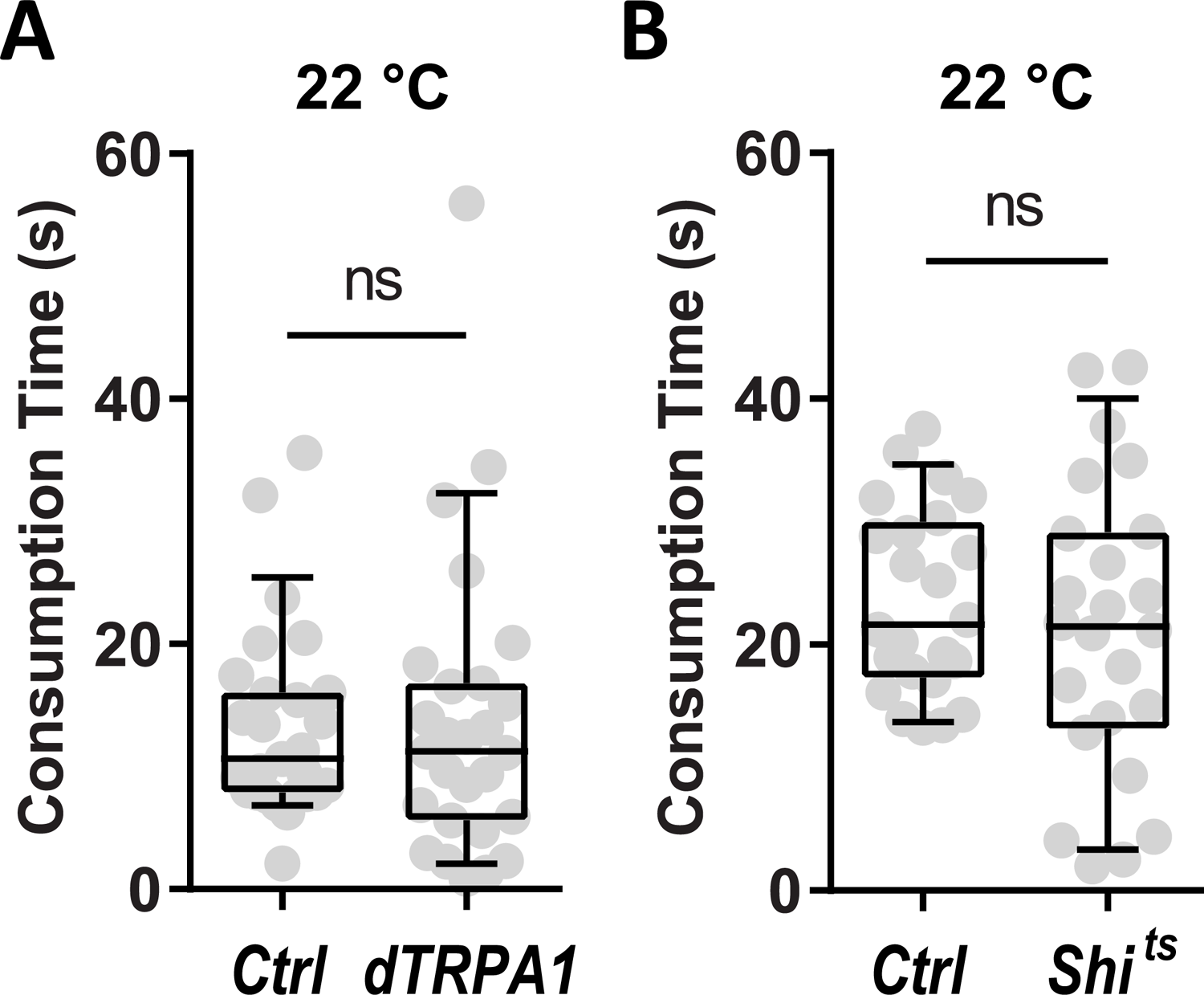
dTRPA1 and Shi^ts^ Expression in Sugar-SELs Did Not Affect Consumption at Permissive Temperature, Related to Figure 4. Controls for Figures 4A-4D. At 22°C when dTRPA1 and Shi^ts^ were not active, their expression in sugar-SELs had no effect on total sucrose consumption time. For all box plots, whiskers = 10^th^ to 90^th^ percentile, box = 25^th^ to 75^th^ percentile, and line in box = median. n = 27 flies/genotype for (A) and n = 24 flies/genotype for (B); Mann Whitney test, ns, not significant.

**Figure S4.**
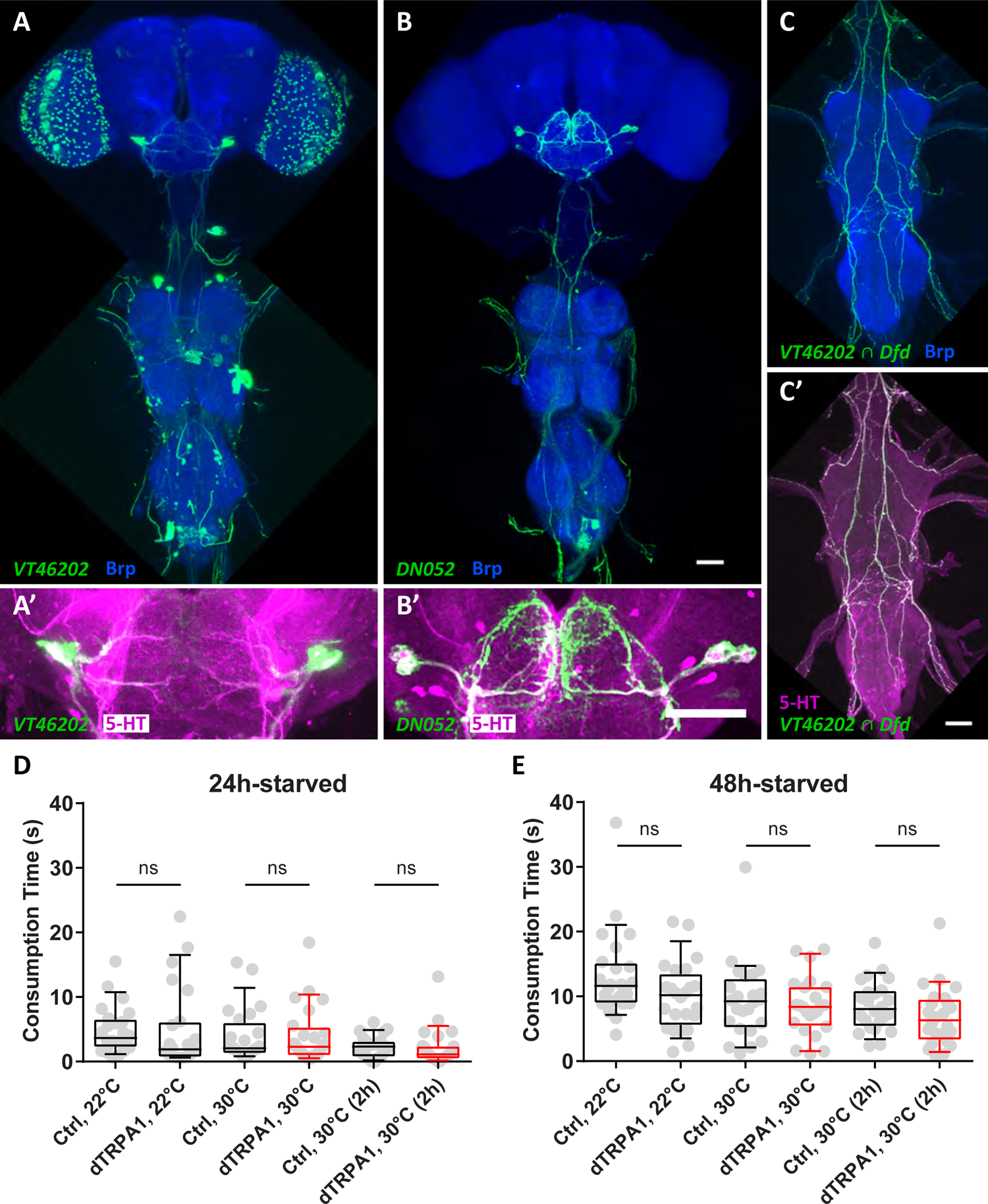
Additional Anatomical and Behavioral Characterization of Bitter-SELs, Related to Figure 5. (A and A’) *VT46202-Gal4* labels bitter-SELs and has sparse expression in the central brain (green). Anti-Brp staining (blue) and anti-5-HT staining (magenta). (B and B’) *DN052 split-Gal4* labels bitter-SELs and a few other cells (green). Anti-Brp staining (blue) and anti-5-HT staining (magenta). Scale bars, 50 μm. (C and C’) Bitter-SEL processes in the VNC in a dorsal-up view (*VT46202 ∩ Dfd*, green). Anti-Brp staining (blue) and anti-5-HT staining (magenta). Scale bar, 50 μm. (D and E) Transient (dTRPA1, 30°C) or prolonged excitation of bitter-SELs for two hours (dTRPA1, 30°C (2h)) had no measurable effect on total sucrose consumption time for flies pre-starved for 24 hours (D) or 48 hours (E). The heat-activated cation channel dTRPA1 is active at 30°C and not active at 22°C. n = 24 flies/genotype; Mann Whitney test, ns, not significant.

**Figure S5.**
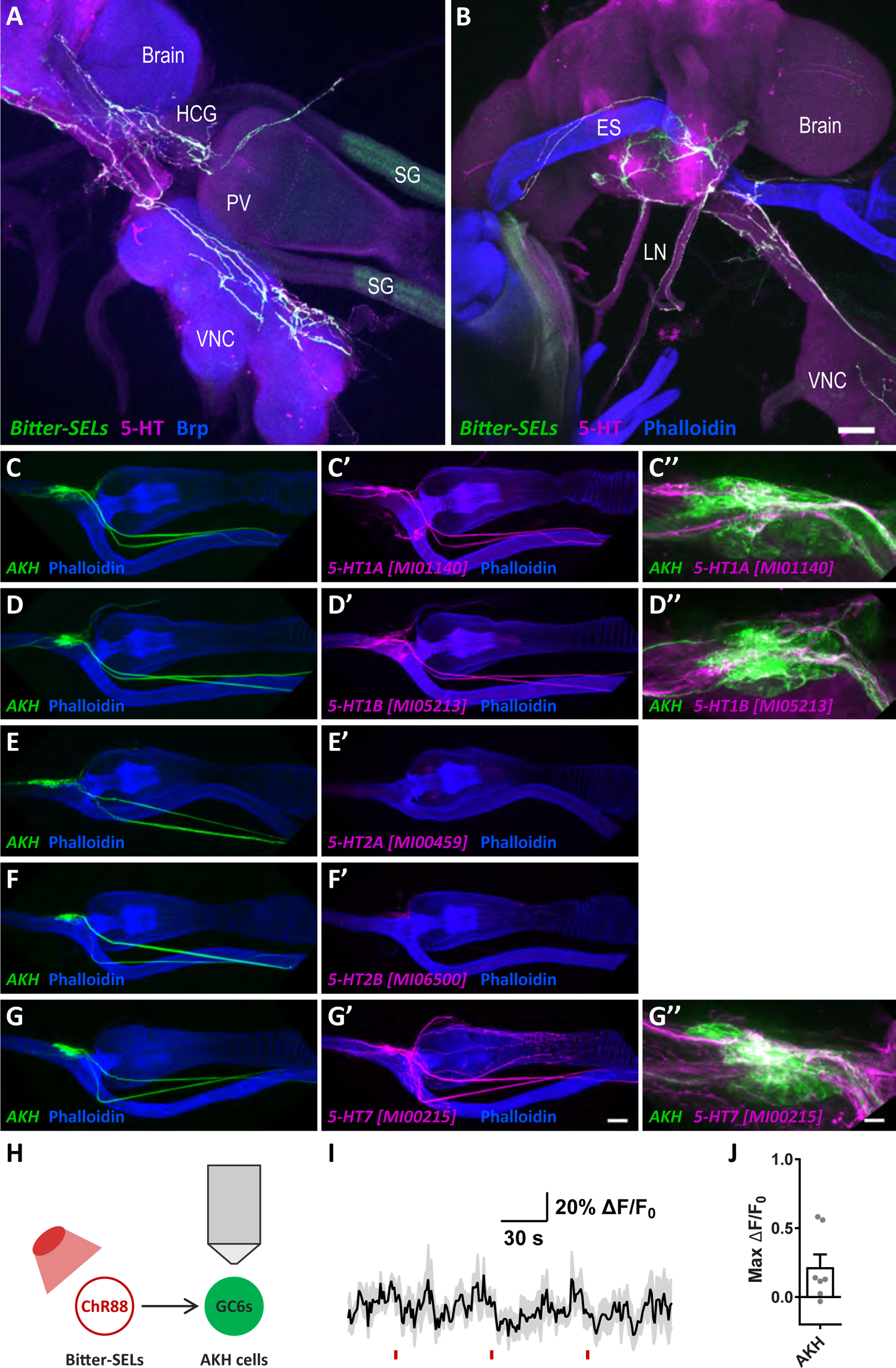
Additional Analysis on Bitter-SELs, AKH Cells, and 5-HT Receptor Expression in the Hypocerebral Ganglion, Related to Figure 6. (A and B) Representative images showing bitter-SEL processes (green) in the central nervous system and the digestive tract. Anti-5-HT staining (magenta); Anti-Brp (blue) labels the central nervous system (A) and phalloidin (blue) labels muscles and the digestive tract (B). VNC, ventral nerve cord; HCG: hypocerebral ganglion; PV, proventriculus; SG, salivary gland; ES, esophagus; LN, labial nerves. Scale bar, 50 μm. (C-G’’) Representative images showing the co-labelling of AKH cells (*AKH-LexA>LexAop-CD8-GFP*, green) and the indicated 5-HT receptors (*5-HT receptor Gal4>UAS-CD8-RFP*, magenta). Phalloidin stains the digestive tract (blue). (C’’), (D’’), and (G’’) are magnified views of the HCG region. Scale bar is 50 μm in (G’) and 10 μm in (G’’). (H-J) Optogenetic excitation of bitter-SELs does not consistently activate AKH cells. (H) Experiment schematic. Chrimson88 (ChR88) was expressed in bitter-SELs while GCaMP6s (GC6s) was independently expressed in AKH cells. (I) Mean GCaMP6s ΔF/F_0_ traces (black lines) ± SEM (gray shading) of AKH cells in response to optogenetic excitation of bitter-SELs (red bars). (J) Scatter plots of maximum ΔF/F_0_ changes for individual flies (n = 7 flies), with bar plots overlay, mean ± SEM.

**Table S1:**
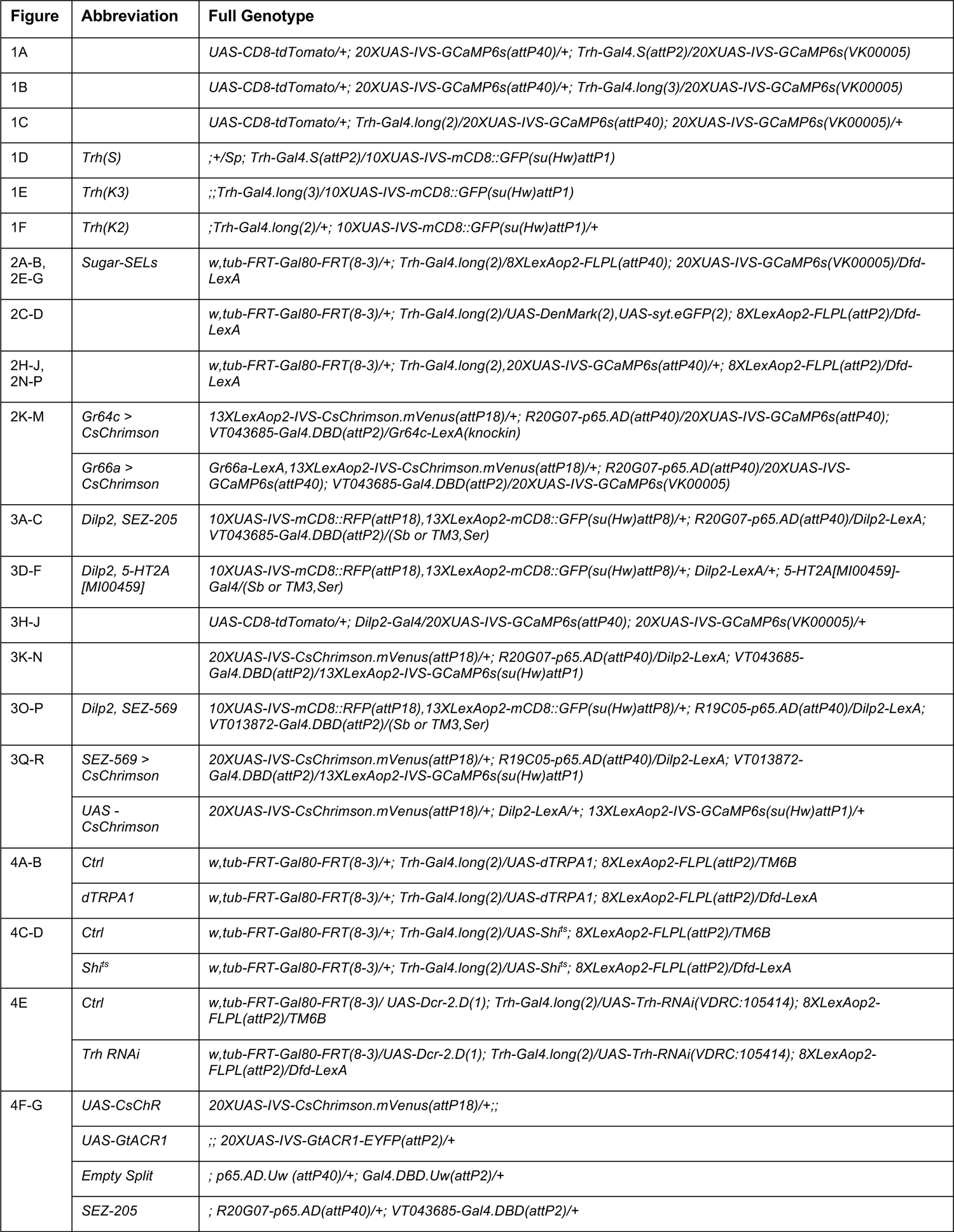
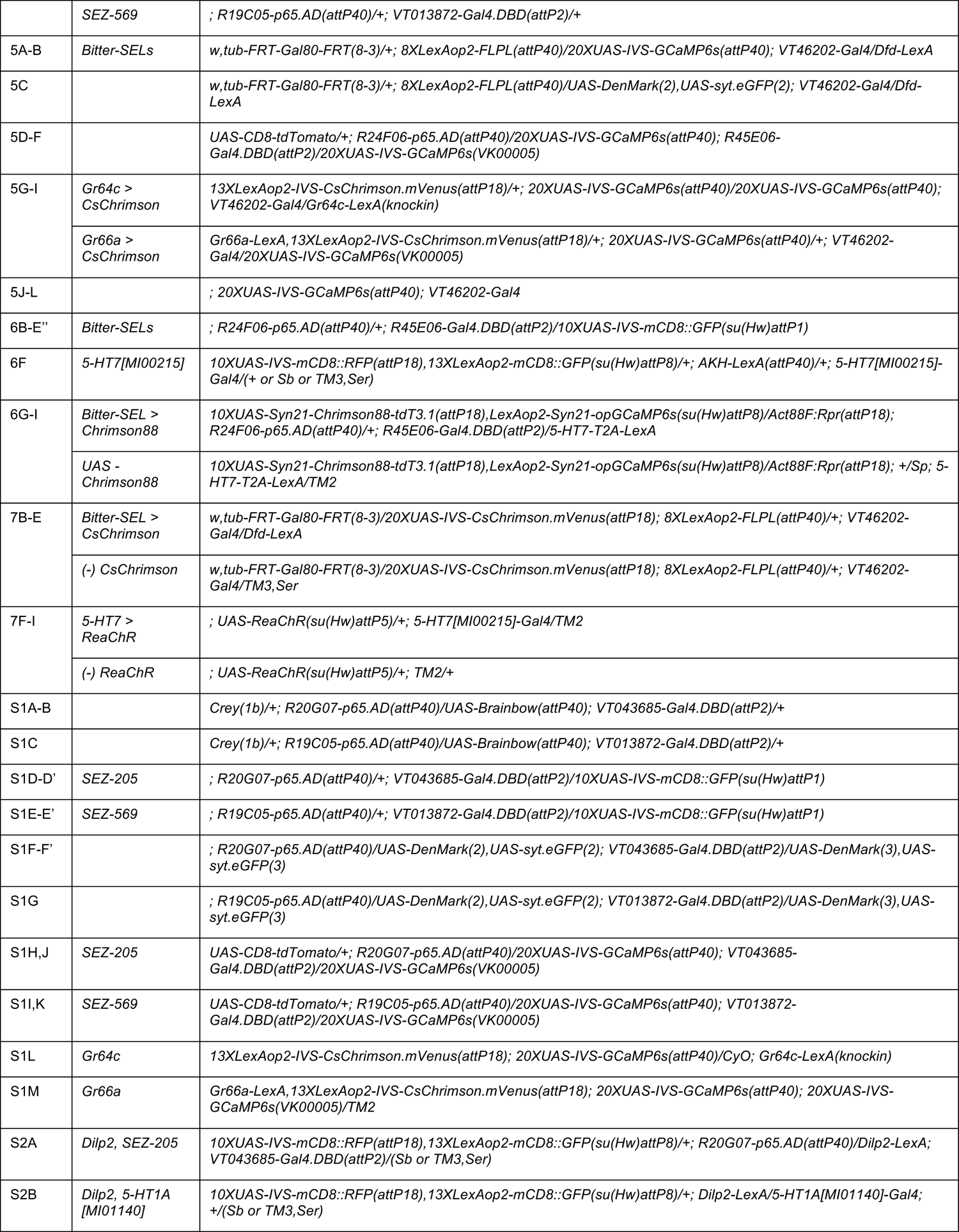
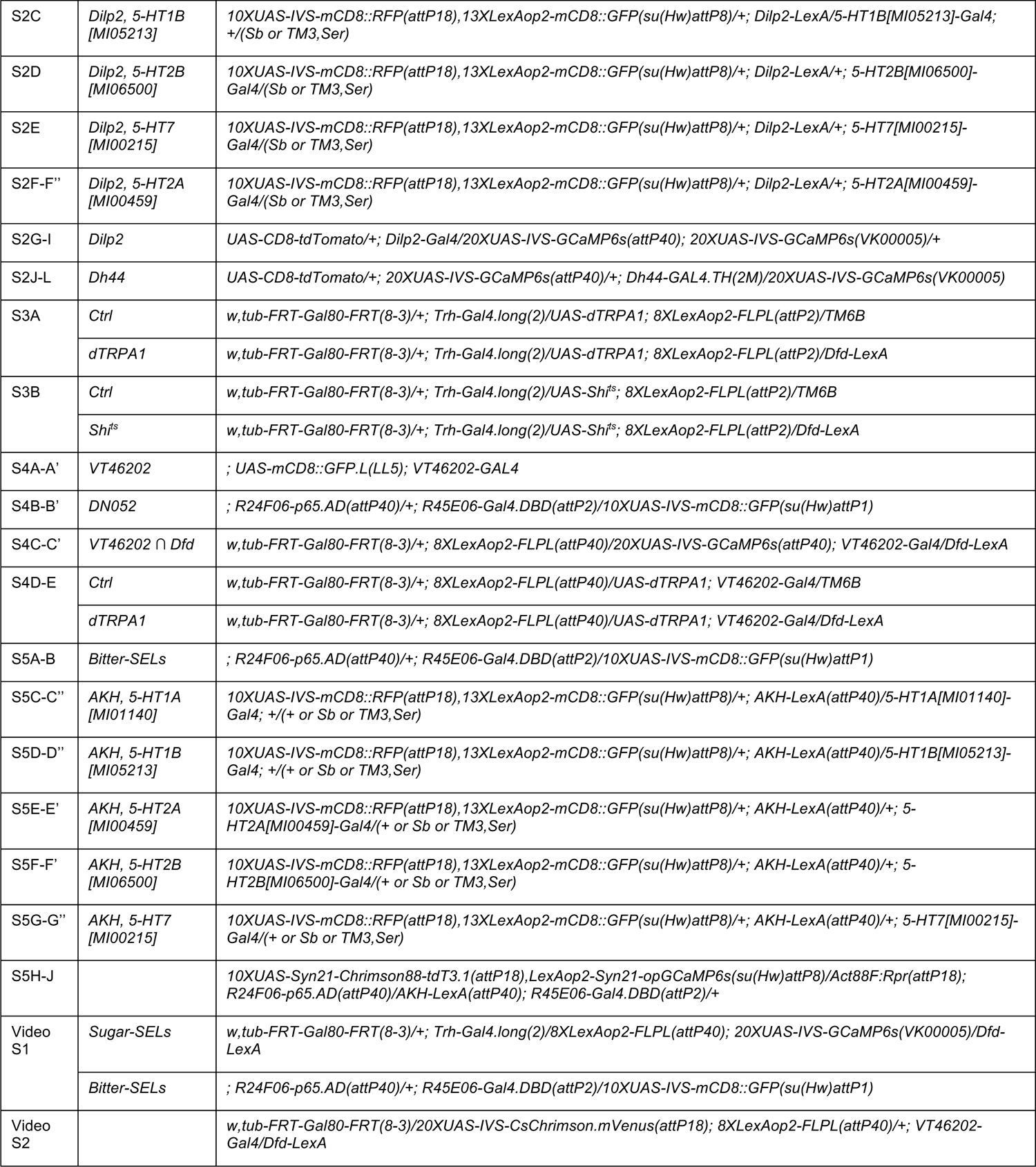
Transgenic Flies Used in This Study, Related to STAR Methods Experimental Model and Subject Details.

**Video S1.** Anatomy of Sugar-SELs and Bitter-SELs in 3D, Related to Figures 2 and 5.

**Video S2.** Example of Crop Contractions Induced by Bitter-SEL Excitation, Related to Figure 7.

## KEY RESOURCES TABLE

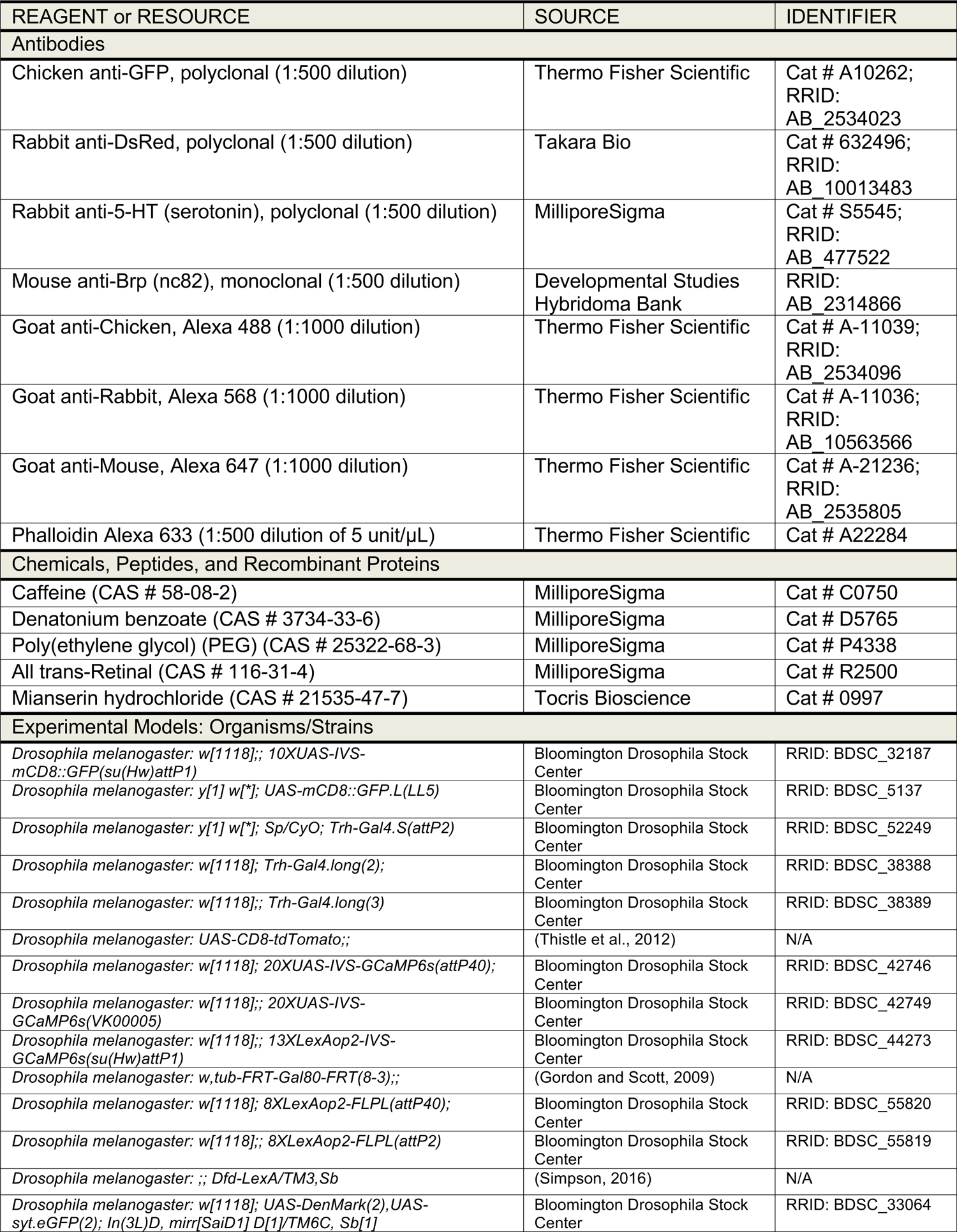
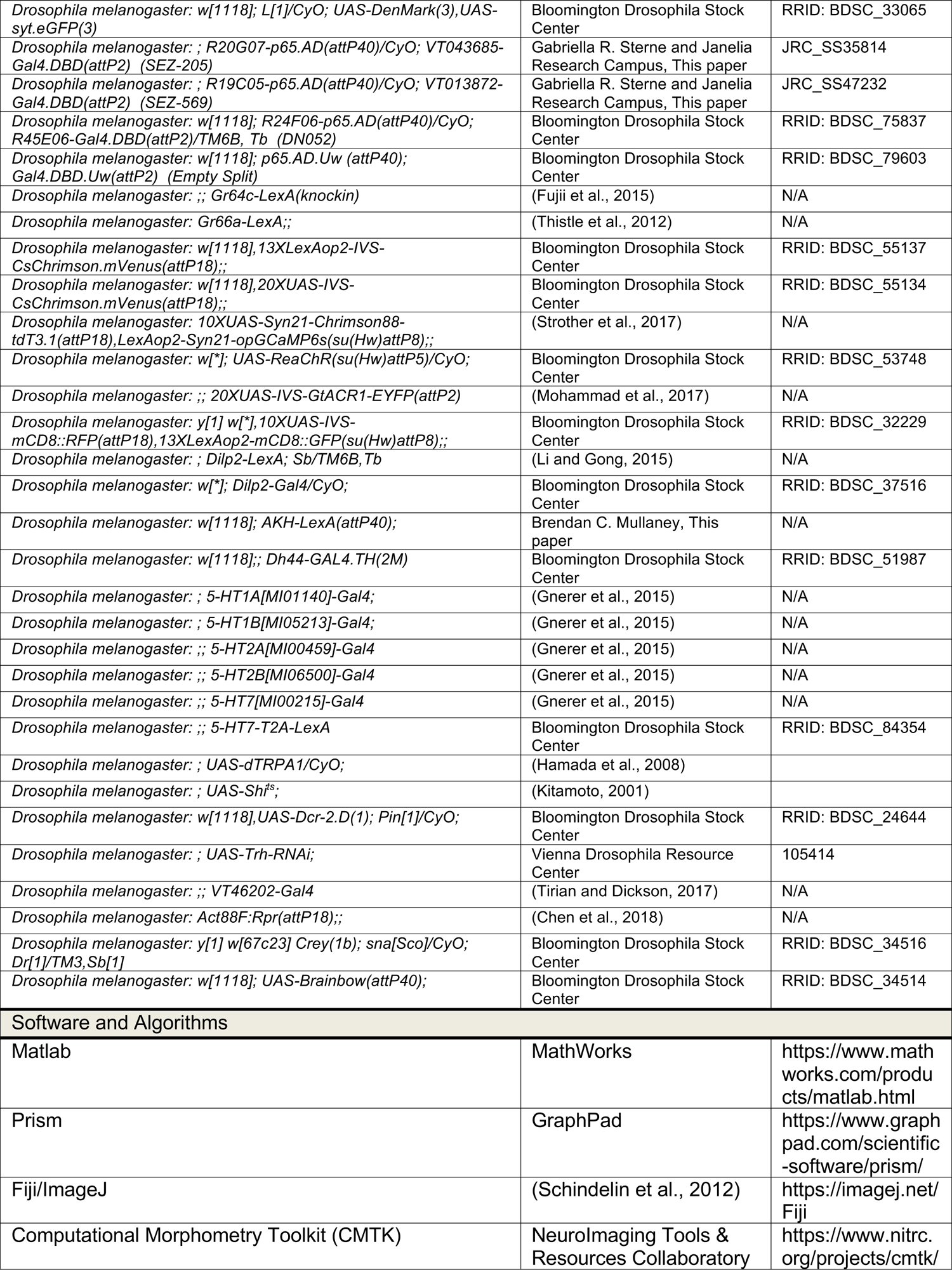
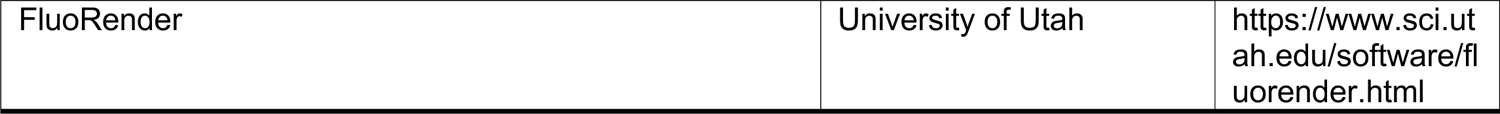

## STAR METHODS

### RESOURCE AVAILABILITY

#### Lead Contact

Further information and requests for resources and reagents should be directed to and will be fulfilled by the Lead Contact, Kristin Scott.

#### Materials Availability

All materials generated in this study including fly lines and custom apparatus will be available upon request.

#### Data and Code Availability

Raw and processed data along with custom analysis scripts will be made available upon request.

### EXPERIMENTAL MODEL AND SUBJECT DETAILS

All experiments were performed in the fruit fly *Drosophila melanogaster*. The Key Resources Table lists the transgenic lines used in this study and Table S1 documents the genotypes used for each figure. Flies were reared on standard cornmeal-yeast-molasses media at 25°C with 65% humidity and a 12hr: 12hr light: dark cycle unless stated otherwise. Flies for optogenetic experiments were raised on standard food and in darkness. Upon eclosion, adult flies were collected and maintained on standard food supplemented with 400 µM all-trans-retinal and in darkness prior to experiments. Flies for dTRPA1, Shi^ts^, and Brainbow experiments were raised at 20-22°C. Adult mated female flies were used for all experiments.

### METHOD DETAILS

#### Genetic Intersections

We used two different intersectional genetic approaches to refine driver expression. The first approach was a genetic intersection between a *Gal4* driver (Brand and Perrimon, 1993; Fischer et al., 1988) and a *LexA* driver (Lai and Lee, 2006). A ubiquitously expressed *Gal80* flanked by two flippase recognition target (*FRT*) sites (i.e., *tub-FRT-Gal80-FRT* (Gordon and Scott, 2009)) was used to suppress *Gal4/UAS* expression. A *LexA* driver was used to express flippase (FLP), which excised the *Gal80* hence allowing *Gal4/UAS* expression. As a result, only cells expressing both *Gal4* and *LexA* expressed the *UAS* transgene. The second approach was the *split Gal4* strategy (Luan et al., 2006). The activation domain of p65 (p65.AD) and the DNA-binding domain of Gal4 (Gal4.DBD) were driven by two different enhancers (Pfeiffer et al., 2010). A functional Gal4-p65 hybrid transcription factor was reconstituted only in cells expressing both enhancers.

#### Calcium Imaging with Taste Stimulation

*In vivo* calcium imaging with taste stimulation was performed as previously described (Harris et al., 2015; Kim et al., 2017). Mated female flies, 5-14 day old, were food-deprived on a wet kimwipe for ∼24 hours prior to imaging. An individual fly was briefly anesthetized using CO_2_ and gently placed into a small slit on a custom-built plastic mount at the cervix so that the head was above the plastic mount while the body was beneath it. The head was then immobilized using nail polish, and the proboscis was gently pulled out using a suction glass pipette and waxed at the rostrum and maxillary palps so that it was in an extended position to allow for taste solution delivery. A piece of coverslip was placed at the base of the rostrum at a 45° angle to the plane of the plastic mount to separate the proboscis from the head. The head was then submerged in adult hemolymph-like (AHL) saline containing 108 mM NaCl, 5 mM KCl, 2 mM CaCl_2_, 2 mM MgCl_2_, 4 mM NaHCO_3_, 1 mM NaH_2_PO_4_, 5 mM trehalose, 10 mM sucrose, 5 mM HEPES (pH 7.5) (Wang et al., 2003). For subesophageal zone (SEZ) imaging, the antennae, the head cuticle above the SEZ, and the underlying air sacs were removed using fine forceps. The esophagus was severed to allow better optical access to the SEZ, except for Figures 5J-5L. For imaging of the median bundle and insulin-producing cells (Figures 2H-2J, 3H-3J, and S2G-S2L), additional cuticle was removed to expose the imaged regions. The esophagus was severed for Figures S2G-S2I, and kept intact for Figures 2H-2J, 3H-3J, and S2J-S2L.

All calcium imaging experiments involving taste stimulation were performed on a fixed-stage Intelligent Imaging Innovations (3i) spinning disk confocal microscope equipped with a Zeiss Plan-APOCHROMAT 20x/1.0 water objective at 1.6x or 1.25x optical zoom. The microscope was equipped with a Piezo drive allowing rapid volumetric scanning. GCaMP6s signal was imaged with a 488 nm laser for 8-14 z-planes encompassing the cells of interest (1.0-1.5 µm steps, 50-100 ms exposure for each plane) at an interval of 1.5-2.0 seconds for 30-60 time-points, using the SlideBook6 acquisition software. Taste stimulation to the proboscis was delivered using a glass capillary (1.0 mm OD/ 0.58 mm ID) filled with ∼5 µL of taste solution. The solution was drawn up the capillary slightly using suction generated by a 1 mL syringe, so that the tip of the capillary was empty. The capillary was then placed over the proboscis using a micromanipulator, visualized under a USB microscope camera. Slight pressure was applied to the syringe to deliver taste solution to the proboscis at desired time-points. To deliver taste stimulation to the legs, a 200 µL pipette tip was attached to a 1 mL syringe and filled with taste solution, and these were secured horizontally on a micromanipulator. A drop of taste solution (∼10 µL) was suspended on the tip of the pipette tip, which was cut to wedge-shaped, and placed in front of the fly prior to image acquisition. The taste solution droplet was manually advanced to contact the legs of the fly at desired time-points, visualized under a USB microscope camera. Sugar solution contained 1 M sucrose, and bitter solution contained 100 mM caffeine, 10 mM denatonium, and 20% polyethylene glycol (PEG). For experiments only involving proboscis taste stimulation (Figures 1A-1C, 2E-2J, 3H-3J, 5D-5F, S1H-S1K, and S2G-S2L), legs were removed. For experiments involving both leg and proboscis taste stimulations (Figures 2N-2P, and 5J-5L), legs were kept intact. For experiments with mianserin (Figures 3H-3J), the insulin-producing cells were first imaged in AHL saline, then AHL saline was replaced by AHL saline supplemented with 100 µM mianserin hydrochloride, and the insulin-producing cells were re-imaged ∼10 minutes later.

#### Calcium Imaging with Optogenetics

*In vivo* calcium imaging with optogenetic stimulation was performed on a Zeiss LSM 880 NLO AxioExaminer equipped with a Coherent Chameleon laser and a Zeiss Plan-APOCHROMAT 20x/1.0 water objective. For imaging of sugar-SELs, bitter-SELs, and insulin-producing cells (Figures 2K-2M, 3K-3N, 3Q, 3R, and 5G-5I), flies were prepared for imaging as described above, except that flies were not food-deprived and their legs were intact. For imaging of 5-HT7(+) enteric neurons and AKH cells (Figures 6G-6I and S5H-S5J), flies were mounted on a recording chamber as described in (Murthy and Turner, 2013), with the dorsal head and thorax accessible for dissection and imaging. The transgene *Act88F:Rpr* in these flies (see Table S1) ablated the indirect flight muscles that fill most of the thorax (Chen et al., 2018), allowing direct optical access to the proventricular region after the dorsal thoracic cuticle and the underlying air sacs and residual muscles were removed using fine forceps. GCaMP6s was imaged on a single plane at 1 Hz with 920 nm 2-photon excitation except for that of the 5-HT7(+) enteric neurons (Figures 6G-6I), which was imaged with a 488 nm laser (at very low intensity, 0.15-0.20%) in order to obtain a thicker z-section to minimize movement artifacts. Optogenetic stimulation was delivered by a mercury lamp filtered at 650 nm (as described in (Bidaye et al., 2020)) or a custom-made 660 nm red diode laser controlled by a pulse generator for 2 s ON and 60 s OFF for three cycles. We used *Gr64c-LexA* to drive *CsChrimson* expression in the sugar GRNs (Figures 2K-2M and 5G-5I) because it does not drive expression in the antennal lobes, in contrast to *Gr5a-LexA* and *Gr64f-LexA* (Figure S1L) (Fujii et al., 2015). Mianserin was applied as described above.

#### Immunohistochemistry

Antibody staining of whole-mount *Drosophila* brain and other tissues was performed as previously described (Yao and Shafer, 2014) with minor modifications. Fly heads (for brain-only staining) or whole flies, with cuticles gently torn open using forceps, were fixed in 3% paraformaldehyde in phosphate buffered saline (PBS) for 1 hr at room temperature. After three washes in PBS, tissues of interest (brain, brain + ventral nerve cord (VNC), guts, or brain + VNC + guts) were dissected in PBS then transferred to PBST (PBS with 0.3% Triton X-100). Tissues were blocked with 5% normal goat serum in PBST for 1 hr at room temperature, then incubated with primary antibodies in block solution at 4 °C for 2-3 days. After five 15-min washes in PBST, tissues were incubated with secondary antibodies in block solution at 4 °C for 1-2 days. After five 15-min washes in PBST, followed by 1-2 exchanges of PBS, tissues were mounted on poly-L-lysine-coated coverslips in PBS and dehydrated in a graded glycerol series (30%, 50%, and 70% glycerol in PBS for 5 min each). The final glycerol solution was replaced with Vectashield Antifade Mounting Medium (H-1000) for imaging and storage. The stained tissues were imaged under a Zeiss LSM 780 or LSM 880 AxioExaminer equipped with a Zeiss Plan-APOCHROMAT 20x/1.0 water objective, using excitation and emission wavelengths corresponding to those of the fluorophores conjugated to the secondary antibodies. Image brightness and contrast were adjusted using Fiji/ImageJ, and image stitching was performed using the Pairwise Stitching plugin in Fiji/ImageJ. Antibodies and their dilutions are described in the Key Resources Table.

#### Analysis of Single-Cell Morphology

We used the *Drosophila* Brainbow technique (Hampel et al., 2011) to analyze the single-cell morphology of sugar-SELs. We initially analyzed the morphology of single sugar-SELs using the genetic intersection of *Trh-Gal4.long(2)* and *Dfd-LexA*. We repeated the analysis using *split Gal4s* for the sugar-SEL PNs (*SEZ-205*) and sugar-SEL LNs (*SEZ-569*) and co-stained for 5-HT (Figures S1A-S1C and Table S1). Flies were raised at 20-22°C without heat shock (because *Crey(1b)* expressed Cre recombinase constitutively), and brains from 7-11 day old mated females were dissected for immunohistochemistry for single-cell morphology analysis.

#### Image Registration to Template Brain

Confocal image stacks of sugar-SELs and bitter-SELs in separate brains were aligned to the brain template JFRC2 (available at https://github.com/jefferislab/BridgingRegistrations) using the anti-Brp (nc82) staining as a reference channel, by nonrigid warping using the Computational Morphometry Toolkit (CMTK) (https://www.nitrc.org/projects/cmtk/), as described in detail in (Jefferis et al., 2007). The registered sugar-SEL and bitter-SEL images, and the JFRC2 brain template were visualized and 3D rendered using FluoRender (https://www.sci.utah.edu/software/fluorender.html) (Video S1).

#### Temporal Consumption Assay

Temporal consumption assay was performed similarly to previously described (Pool et al., 2014). Adult mated female flies, 7-15 day old, were food-deprived on a wet kimwipe for ∼24 hours (unless stated otherwise). Flies were mounted on glass slides with nail polish and allowed to recover in a humidified chamber for ∼2-3 hours. Individual flies were presented with a drop of 1 M sucrose (supplemented with 0.25 mg/mL FD&C No. 1 blue dye for visualization) from a 200 µL pipette tip attached to a 1 mL syringe and allowed to consume for at least 10 times and until the flies no longer consumed.

Feeding bouts were video recorded at 30 frames per second using a USB microscope camera and manually annotated (Figures 4A-4E and S3) or manually recorded using an online chronometer (http://online-stopwatch.chronme.com/) (Figures 4F-4G, S4D, and S4E). Flies for dTRPA1 and Shi^ts^ experiments were raised 20-22°C. To activate dTRPA1, flies were placed on a 29-30°C heat block for at least 5 min before testing and throughout testing. For Shi^ts^ silencing experiments, flies were placed in a humidified chamber at 30-32°C for ∼2-3 hours and then tested on a 30-32°C heat block. Sibling controls from the same genetic crosses were used for both sets of experiments. Flies for CsChrimson and GtACR1 experiments were raised on standard food at 25°C in darkness. Upon eclosion, adult flies were collected and maintained on standard food supplemented with 400 µM all-trans-retinal until they were food-deprived for the consumption assay. CsChrimson was activated by a Laserglow red laser (635-650 nm) and GtACR1 was activated by a Laserglow green laser (532 nm) for ∼1 min before testing and throughout testing.

#### Crop Contraction Assay

The crop contraction assay was modified from (Solari et al., 2017). We used flies fed *ad libitum* on fresh food for 3-5 days because they typically had few basal crop contractions. A live fly was briefly anesthetized with ice and mounted on the bottom of a 35 mm petri dish using petroleum jelly with its ventral side facing up. The legs were removed, the proboscis was sealed with wax, and AHL saline was added to the dish to submerge the fly. The cuticle in the ventral upper abdomen was torn open using fine forceps with care not to damage the crop or gut tissues, exposing the crop for video recording. The above procedures were carried out at low light intensity, and the fly was allowed to recover in darkness for ∼4 min before recording began. Crop contractions were video recorded under 850 nm infrared light illumination at 10 frames per second, using a FLIR Blackfly S USB3 camera (BFS-U3-13Y3M) mounted on an Olympus SZX16 stereo microscope. Optogenetic stimulation was delivered by a custom-made 630 nm LED panel controlled by a pulse generator (100 Hz of 5 ms pulses for 30 s). A 760 nm longpass filter was fitted in front of the camera to prevent the 630 nm stimulation light from interfering with video recording (Figure 7A).

### QUANTIFICATION AND STATISTICAL ANALYSIS

All statistical tests were performed in Prism. Data was tested for normal distribution using the D’Agostino & Pearson normality test. In general, if all data for comparison passed the normality test (alpha = 0.05), parametric tests were used; otherwise, nonparametric tests were used. For similar experiments, the same statistical tests were performed for consistency.

#### Calcium Imaging with Taste Stimulation

The volumetric GCaMP6s images for taste stimulation experiments were collapsed in the z-axis to generate a max-intensity z-projection image (referred to as max-z image hereafter) for each time-point. This max-z image sequence was corrected for movement artifacts using the StackReg plugin in Fiji/ImageJ with ‘Rigid Body’ or ‘Translation’ transformation. The movement-corrected max-z image sequence was used for subsequent analyses.

To generate a ΔF/F_0_ image, GCaMP6s images (typically of four time-points) before stimulation were averaged to generate a baseline F_0_ image. GCaMP6s images (typically of three time-points) during peak response to a stimulation were max-intensity projected to generate an F_max_ image. The ΔF/F_0_ image was calculated as (F_max_ − F_0_) / F_0_ for each pixel. The only exception is the ΔF/F_0_ images for *Trh-Gal4s* (Figures 1A-1C). Because of the dense GCaMP6s-expressing arbors, movement correction was performed for each z-plane over time and a ΔF/F_0_ image was generated for each z-plane. The ΔF/F_0_ images for all z-plane were max-intensity projected to generate the final ΔF/F_0_ images shown in Figures 1A-1C. Image calculations were done in Matlab and ΔF/F_0_ images were displayed using Fiji/ImageJ.

To generate ΔF/F_0_ traces, regions of interest (ROIs) were manually drawn on GCaMP6s-expressing processes in Fiji/ImageJ, and the average fluorescence intensity of each ROI over time was measured using the Time Series Analyzer V3 plugin in Fiji/ImageJ. A large background ROI was drawn on areas without GCaMP6s expression, and the average fluorescence intensity of the background ROI for each time-point was subtracted from that of each ROI to generate a background-corrected ROI fluorescence intensity over time, F(t). For each ROI F(t) trace, fluorescence intensity (typically of four time-points) before stimulation was averaged to generate a fluorescence baseline F_0_, and ΔF/F_0_ was calculated as (F(t) − F_0_) / F_0_ for each time-point (t). If multiple ROIs were drawn for a single cell/cell type, the ΔF/F_0_ traces for those ROIs were averaged to generate a mean ΔF/F_0_ trace for that cell/cell type. Max ΔF/F_0_ was the maximum ΔF/F_0_ value during and immediately after stimulation for a given ΔF/F_0_ trace. ROI drawing and intensity measures were done in Fiji/ImageJ. Intensity calculations were done in Matlab. Statistical tests were performed in Prism and reported in the figure legends.

#### Calcium Imaging with Optogenetics

The analyses of GCaMP6s imaging data for optogenetic experiments were performed in the same way as those described for the taste stimulation experiments except for the following: (1) Because only a single plane was imaged for the optogenetic experiments, max-intensity z-projection was not performed. (2) Typically, five time-points before stimulation were used to generate the F_0_ images and F_0_, and three to four time-points during peak response to stimulation were used to generate the F_max_ images. (3) The background fluorescence levels were very low in these images and often resulted in erroneously large pixel values when ΔF/F_0_ images were calculated. To minimize such errors, an intensity-based threshold was applied to the image sequence to exclude areas without GCaMP6s expression from calculations.

#### Temporal Consumption Assay

Feeding bouts of representative individuals (Figures 4A and 4C) were plotted in Matlab using the start and end times of each feeding bout. The total consumption time for each individual was the sum of the duration of each feeding bout. Statistical tests were performed in Prism and reported in the figure legends.

#### Crop Contraction Assay

To quantify crop contractions, a region of interest (ROI) was drawn encircling the crop, and the ROI frame-to-frame change of individual pixel intensity was calculated and summed per frame. This total change of pixel intensity per frame was then normalized to the pre-stimulation baseline (the 25th percentile of the 30 s pre-stimulation period) to generate a time-series fold change as shown in Figures 7B and 7F. This quantification method accurately reflected the timing and amplitude of crop contractions as seen visually (Video S2). ROI drawing was done in Fiji/ImageJ and calculations were done in Matlab. Statistical tests were performed in Prism and reported in the figure legends.

